# A cross-species proteomic map reveals neoteny of human synapse development

**DOI:** 10.1101/2022.10.24.513541

**Authors:** Li Wang, Kaifang Pang, Li Zhou, Arantxa Cebrián-Silla, Susana González-Granero, Shaohui Wang, Qiuli Bi, Matthew L. White, Brandon Ho, Jiani Li, Tao Li, Yonatan Perez, Eric J. Huang, Ethan A. Winkler, Mercedes F. Paredes, Rothem Kovner, Nenad Sestan, Alex A. Pollen, Pengyuan Liu, Jingjing Li, Xianhua Piao, José Manuel García-Verdugo, Arturo Alvarez-Buylla, Zhandong Liu, Arnold R. Kriegstein

## Abstract

The molecular mechanisms and evolutionary changes accompanying synapse development are still poorly understood. Here, we generated a cross-species proteomic map of synapse development in the human, macaque, and mouse neocortex. By tracking the changes of >1,000 postsynaptic density (PSD) proteins from midgestation to young adulthood, we found that PSD maturation in humans separates into three major phases that are dominated by distinct pathways. Cross-species comparisons reveal that the human PSD matures about two to three times slower than other species and contains higher levels of Rho guanine nucleotide exchange factors (RhoGEFs) in the perinatal period. Enhancement of the RhoGEF signaling in human neurons delays the morphological maturation of dendritic spines and the functional maturation of synapses, potentially contributing to the neotenic traits of human brain development. In addition, PSD proteins can be divided into four modules that exert stage- and cell type-specific functions, possibly explaining their differential associations with cognitive functions and diseases. Together, our proteomic map of synapse development provides a blueprint for studying the molecular basis and evolutionary changes of synapse maturation.

## Main Text

Synapses establish the neuronal networks that mediate information processing in the brain. Synaptic dysfunction plays a critical role in most brain diseases, including disorders that typically occur in childhood, adolescence, or adulthood ^1–3^. Therefore, understanding synapse formation, maturation, and specification is essential for understanding human cognition and mental disorders.

The two major types of chemical synapses in the brain are excitatory glutamatergic synapses and inhibitory GABAergic synapses. They differ in neurotransmitters, morphology, molecular composition, and postsynaptic organization ^4–7^. The postsynaptic density (PSD) of excitatory synapses is a highly specialized structure located beneath the postsynaptic membrane and is more prominent than its counterpart at inhibitory synapses. Biochemical isolation of PSDs from the adult brain followed by mass spectrometry revealed that the PSD is a highly sophisticated protein complex composed of >1,000 proteins including cytoskeletal proteins, neurotransmitter receptors, signaling enzymes, ribosomal proteins, and scaffolding proteins ^8^. Mutations in these proteins cause over 130 brain diseases ^9^.

Excitatory synapses and associated PSDs undergo profound changes at both morphological and compositional levels during brain development ^10–14^. In particular, developmental increases in the ratio of the N-methyl-D-aspartic acid (NMDA) receptor subunits GRIN2A to GRIN2B and of the PSD scaffolding proteins DLG4 to DLG3 are critical for the functional maturation of synapses ^15–,17^. However, studies to understand the developmental changes of the PSD are limited to dozens of well-known PSD proteins typically identified in the adult brain ^10,12,13^. Unbiased, systematic characterization has been limited, especially in humans ^18^. In addition, synapse density, composition, and maturation rates differ between species, potentially contributing to the evolutionary variation of neurotransmission properties, cognitive ability, and behavioral repertoire ^19–25^. For example, prolonged maturation or neoteny of human synapses has been suggested as a possible explanation for the emergence of human-specific cognitive traits ^26–28^. Nevertheless, we still know little about the underlying molecular mechanisms.

Here, we generate a cross-species proteomic map of synapse development in the neocortex, identifying the dynamics of >1,000 PSD proteins and the molecular pathways that govern individual phases of synapse maturation. A comparison of the maturing PSDs in humans to those in macaques and mice reveals that PSD maturation in humans is two to three times slower than that in other species. Moreover, Rho guanine nucleotide exchange factors (RhoGEFs), which serve to delay synapse maturation, are more abundant in human PSDs in the perinatal phase, possibly contributing to the neotenic traits of human synapses. Integrating these data with transcriptomic and genetic data, we further determine the gene regulatory network, cell type specificity, and selective disease vulnerability of synapse maturation. Our data provide a temporal map of the topology of synapse development in the neocortex and offer insight into the evolutionary mechanisms of synaptic neoteny in humans.

## Results

### Changes in PSD composition during human neocortical development

To understand the molecular changes of the PSD in the developing human neocortex, we obtained neocortical samples across six major developmental stages ranging from the second trimester to young adulthood (Fig. 1a, Supplementary Table 1). The six stages were chosen to cover major developmental events including neurogenesis, neuronal migration, synaptogenesis, myelination, and synaptic pruning. We used samples from the prefrontal cortex (PFC) to reduce the confounding effect of cortical areas, except for second trimester samples that lacked area information due to limited availability. PSDs were isolated from each sample as described previously ^29^. Isolation of the PSD, including from immature human brain samples, was successful as indicated by the following quality control metrics. Integral components of the PSD, but not presynaptic (SYP) or cytoplasmic (GAPDH) proteins, were enriched in the PSD fraction of early-stage samples (Extended Data Fig. 1a). In addition, electron microscopy identified typical PSD-like electron-dense structures in the PSD fraction of immature samples (Extended Data Fig. 1b). Furthermore, GRIN2B and DLG4, two PSD proteins that decrease and increase, respectively, during PSD maturation ^12^, showed the expected temporal abundance patterns in isolated PSDs (Extended Data Fig. 1c). Finally, the yield of PSDs correlated well with the estimated number of synapses (Extended Data Fig. 1d).

**Fig. 1.**
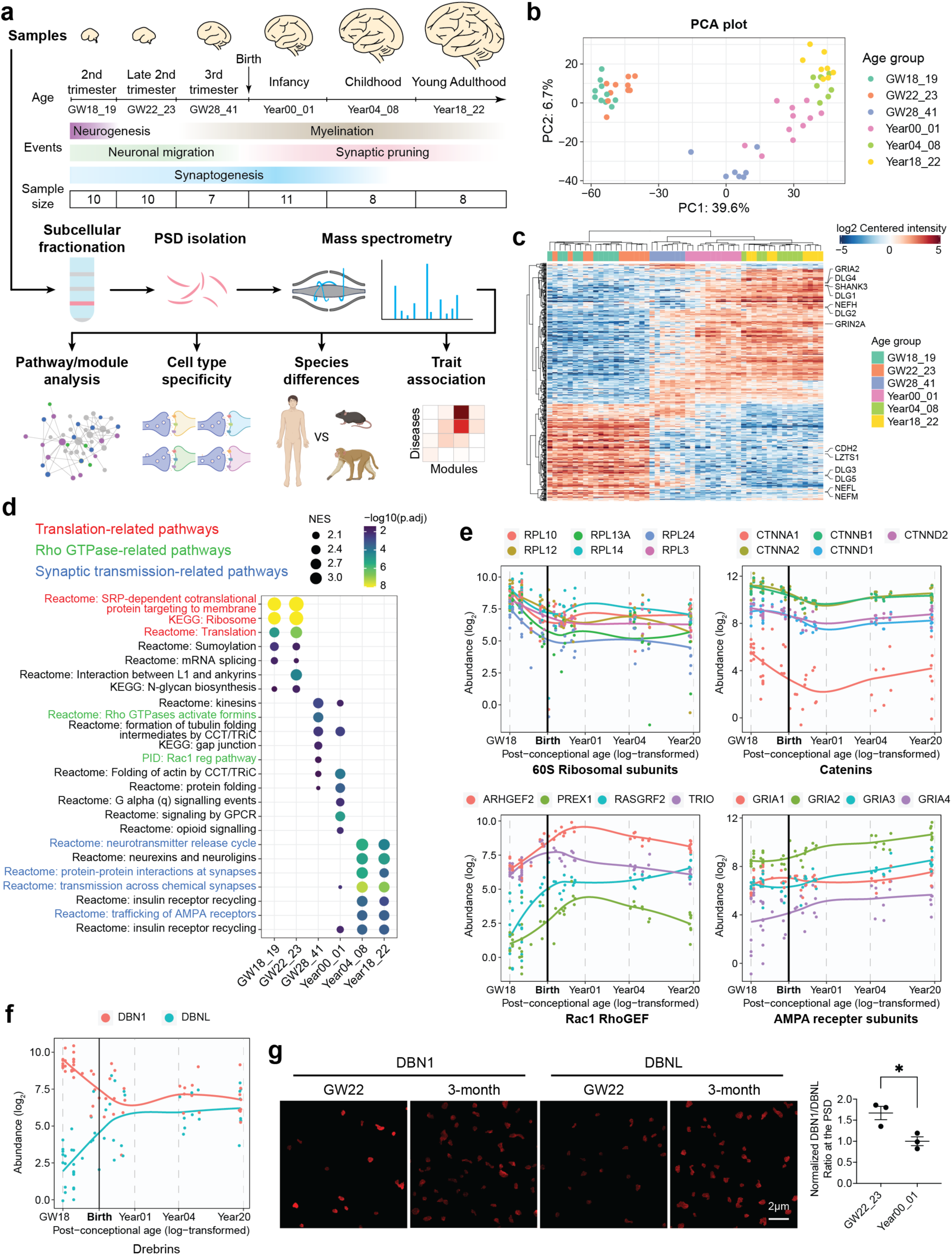
Changes in PSD composition during human neocortical development. **a**, Flow chart of the overall approach. **b**, PCA plots of the samples colored by their age groups. **c**, Hierarchical clustering of the samples based on proteins with differential abundance. **d**, Gene set enrichment analysis for individual age groups. NES: normalized enrichment score. **e**, Abundance patterns of representative PSD proteins. **f**, Abundance patterns of DBN1 and DBNL. **g**, Immunofluorescent intensity of DBN1 and DBNL at DLG4 loci in the human neocortex (n = 3, 3 samples, scale bar: 2 μm). *p < 0.05; unpaired two-tailed *t* test.

We performed liquid chromatography and tandem mass spectrometry (LC-MS/MS) analysis and label-free quantification on 54 PSD samples. Each PSD sample was isolated from a different neurologically normal individual and had passed screening for synaptic proteome preservation ^30^. The identified proteins overlapped significantly with previously reported PSD proteins at comparable stages (Extended Data Fig. 1e) ^9,31^. After removing potential contaminants, we found a total of 1765 PSD proteins with some proteins being stage-specific (Extended Data Fig. 1f, Supplementary Table 2). To assess the quality of the data, we first sought to determine whether developmental changes were the main driver of variance. Principal component (PC) analysis revealed that samples from the same age group were closely clustered (Fig. 1b). PC1, accounting for 39.5% of the variability, was strongly correlated with the age of the samples but not with other potential confounding factors like sex or processing batch (Extended Data Fig. 1g). Moreover, variance across age groups explains a median of 41.7% of the variation in the dataset, after correcting for processing batch, PSD quality, and sex (Extended Data Fig. 1g). Hierarchical clustering also showed that samples were clustered by age (Fig. 1c). Proteins such as GRIN2A, GRIN2B, DLG3, and DLG4 showed the expected abundance patterns during PSD maturation and were consistent with Western blotting data (Fig. 1c, Extended Data Fig. 2a,b). To validate the identified PSD proteins *in situ* in the immature human neocortex, we performed immunostaining of several proteins that show enrichment at midgestation, including the ribosomal subunit RPS6, β-catenin (CTNNB1), the vesicle trafficking regulator GDI1, and the actin modulator cofilin (CFL1). All these proteins colocalized with the canonical PSD marker DLG4 in a subset of synapses (Extended Data Fig. 3).

We performed gene set enrichment analysis (GSEA) to identify molecular pathways with higher activity at individual developmental stages compared with other stages. In general, PSD maturation appears to undergo three major phases (midgestational, perinatal, and postnatal). The midgestational phase, between gestational week 18 to 23, was enriched for translation-related pathways (Fig. 1d,e, Extended Data Fig. 2c). The perinatal phase, between the third trimester and one year of age, was enriched for Rho GTPase and protein folding pathways (Fig. 1d,e, Extended Data Fig. 2d). The postnatal phase, above four years of age, was enriched for synaptic transmission-related pathways and neurexin/neuroligin-associated proteins which play instructive roles in synapse formation and maturation ^32^ (Fig. 1d,e, Extended Data Fig. 2e). These results suggest that local protein synthesis, actin cytoskeleton reorganization, and enhancement of synaptic efficacy were sequentially activated during PSD development. At the individual protein level, proteins from the same complex or pathway generally exhibit similar abundance changes during development (Fig. 1e, Extended Data Fig. 2c–e). However, relative changes in the abundance of homologous proteins such as GRIN2A/GRIN2B and DLG3/DLG4, as shown above, are critical for synapse maturation ^15–17^. Another example concerns the two predominant α-amino-3-hydroxy-5-methyl-4-isoxazolepropionic acid (AMPA) receptor subunits GRIA1 and GRIA2. We found that GRIA2 increased steadily during development, whereas GRIA1 remained relatively constant (Fig. 1e), consistent with GRIA2-lacking, and thus calcium-permeable AMPA receptors being essential for early synaptic function ^33^. In addition to these proteins, we discovered many other homologous proteins that exhibited reciprocal pattern changes (Fig. 1f, Extended Data Fig. 4), including drebrin/drebrin-like proteins, whose abundance changes were further validated by immunostaining (Fig. 1g, Extended Data Fig. 5). These newly identified reciprocal changes may also play important roles in PSD maturation.

### Protein modules and their coordinated functions in the PSD

Proteins with similar abundance patterns during PSD maturation could represent protein modules with specific molecular functions. We identified four protein modules based on correlation by weighted correlation network analysis (WGCNA) (Fig. 2a) ^34^. All four modules were significantly enriched for protein-protein interactions (PPIs), whereas no enrichment was found for proteins with no module assignment (the grey module) (Fig. 2b), suggesting that proteins in the same module work synergistically by forming protein complexes. Indeed, pathway overrepresentation analysis highlighted module-specific enrichment in particular biological pathways (Fig. 2c, Supplementary Table 3). Specifically, the brown, blue, turquoise, and yellow modules, ranked by their timing of peak abundance, were enriched with translation-related pathways, axon guidance-related pathways, Rho GTPase-related pathways, and synaptic transmission-related pathways, respectively (Fig. 2c). Similar results were obtained by synaptic gene ontology (SynGO) enrichment analysis (Extended Data Fig. 6a, Supplementary Table 3) ^35^. In summary, abundance patterns and molecular functions of the PSD protein modules are consistent with the GSEA results at individual developmental stages.

**Fig. 2.**
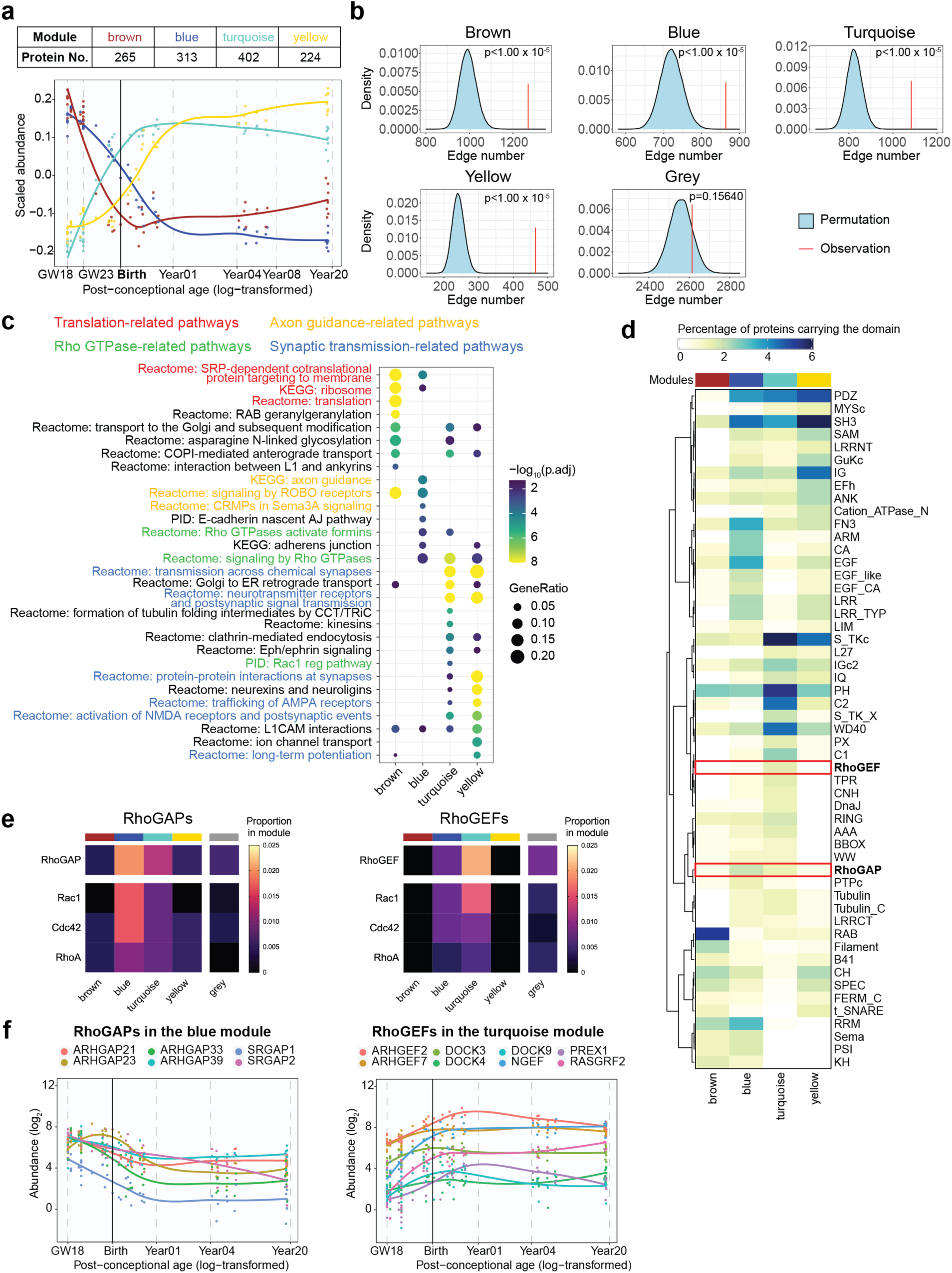
Protein modules of the developing human PSD with distinct functions. **a**, Scaled abundance patterns (module eigengene values) of four protein modules of the human PSD identified by WGCNA. **b**, Kernel density estimation of the null distributions of protein-protein interaction (PPI) numbers assuming no enrichment of PPI in individual modules; the vertical red lines indicate the observed PPI numbers in each module. **c**, Pathway enrichment analysis of each module (hypergeometric test). **d**, Distribution of protein domains in each module. **e**, Proportions of RhoGAPs and RhoGEFs and their subtypes in each module. **f**, Abundance patterns of RhoGAPs in the blue module and RhoGEFs in the turquoise module.

To visualize potential protein complexes and interactions in the PSD, we generated PPI-co-abundance networks in each module (see Methods) (Extended Data Figs. 6b, 7, 8). As expected, proteins involved in the same pathway were clustered more closely, as indicated by a shorter average path length to each other than to proteins outside the pathway (Extended Data Fig. 6b,c). PPIs and biological functions of a protein are often mediated by protein domains. We determined the distribution of protein domains in the modules (Fig. 2d) and the domain architecture of individual proteins (Extended Data Fig. 9, Supplementary Table 4). Domains involved in vesicle trafficking (RAB and t_SNARE), cell adhesion (LRR, CA, and ARM), signal transduction (S_TKc, C1, and C2), and adult PSD scaffolds (PDZ, SH3, and GuKc) ^13^ were enriched in the brown, blue, turquoise, and yellow modules, respectively (Fig. 2d). Interestingly, although both the blue and turquoise modules were involved in Rho GTPase signaling (Fig. 2d), RhoGAP and RhoGEF domains were selectively enriched in each of them (Fig. 2d). Indeed, Rho GTPase-activating proteins (RhoGAPs), particularly those specific for Rac1 and Cdc42, were enriched in the blue module (Fig. 2e,f, Extended Data Fig. 10a–c, Supplementary Table 4). Conversely, Rho guanine nucleotide exchange factors (RhoGEFs), particularly those specific for Rac1, were enriched in the turquoise module (Fig. 2e,f, Extended Data Fig. 10d,e, Supplementary Table 4). Given that RhoGAP and RhoGEF proteins exert antagonistic functions in activating Rho GTPases, these results suggest that synaptic Rho GTPases are gradually shifting towards a more active state during PSD maturation to facilitate stage-specific cytoskeleton reorganization requirements and morphological changes.

### Generalization to other cortical regions

Our initial analysis focused on the PFC. To address whether our findings are applicable to other cortical regions, we conducted a similar analysis on PSD samples from human primary visual cortex (V1), which is located on the opposite pole of the rostral-caudal axis from PFC and is a sensory cortical area (Extended Data Fig. 11a, Supplementary Table 1, Supplementary Table 5). Our analysis found that, like PFC, V1 samples were separated into three clusters corresponding to the midgestational, perinatal, and postnatal phases (Extended Data Fig. 11b,c). Additionally, we observed similar pathway enrichment patterns with translation-, Rho GTPase-, and synaptic transmission-related pathways activated sequentially from midgestation to adulthood (Extended Data Fig. 11d, Supplementary Table 5). Further, the four PSD modules showed similar abundance patterns in V1 (Extended Data Fig. 11e). To compare the PSD samples quantitatively between the two regions, we calculated the Pearson correlation coefficients and found that they correlated well with counterparts in the same phase, with a Pearson r > 0.75 (Extended Data Fig. 11f). Therefore, our results suggest that the major findings from the PFC dataset can be generalized to other regions of the human neocortex, including V1.

### Transcription of PSD proteins and its cell type specificity

To understand the role of transcription in regulating PSD development, we compared the RNA levels of PSD modules with their abundance patterns. Integrating BrainSpan and PsychENCODE transcriptomic data ^36,37^ from the developing human neocortex with our proteomic data, we found that the general trends were preserved. However, the respective differences between the brown/blue and turquoise/yellow modules were greatly diminished (Fig. 3a, Supplementary Table 5). For example, Rho GTPase regulators and PSD scaffolding proteins from the turquoise and yellow modules, respectively, had distinct abundance patterns in the proteomic data, but they had similar expression patterns in the transcriptomic data (Fig. 3b). To quantify the concordance between RNA and protein, we calculated the Spearman’s rank coefficient of correlation between RNA and protein levels of all PSD proteins (Supplementary Table 5). We found that proteins in the blue and yellow modules generally had high RNA-protein concordance (median Spearman r > 0.5) (Fig. 3c). In contrast, brown and turquoise modules had significantly lower concordance (Fig. 3c), suggesting that post-transcriptional/translational regulatory mechanisms such as protein trafficking and turnover play key roles in regulating their PSD abundance. Consistent with these results, module density and connectivity preservation analysis ^38^ showed that although all four modules were at least moderately preserved in the transcriptomic data, the brown and turquoise modules were among the least preserved (Extended Data Fig. 12a).

**Fig. 3.**
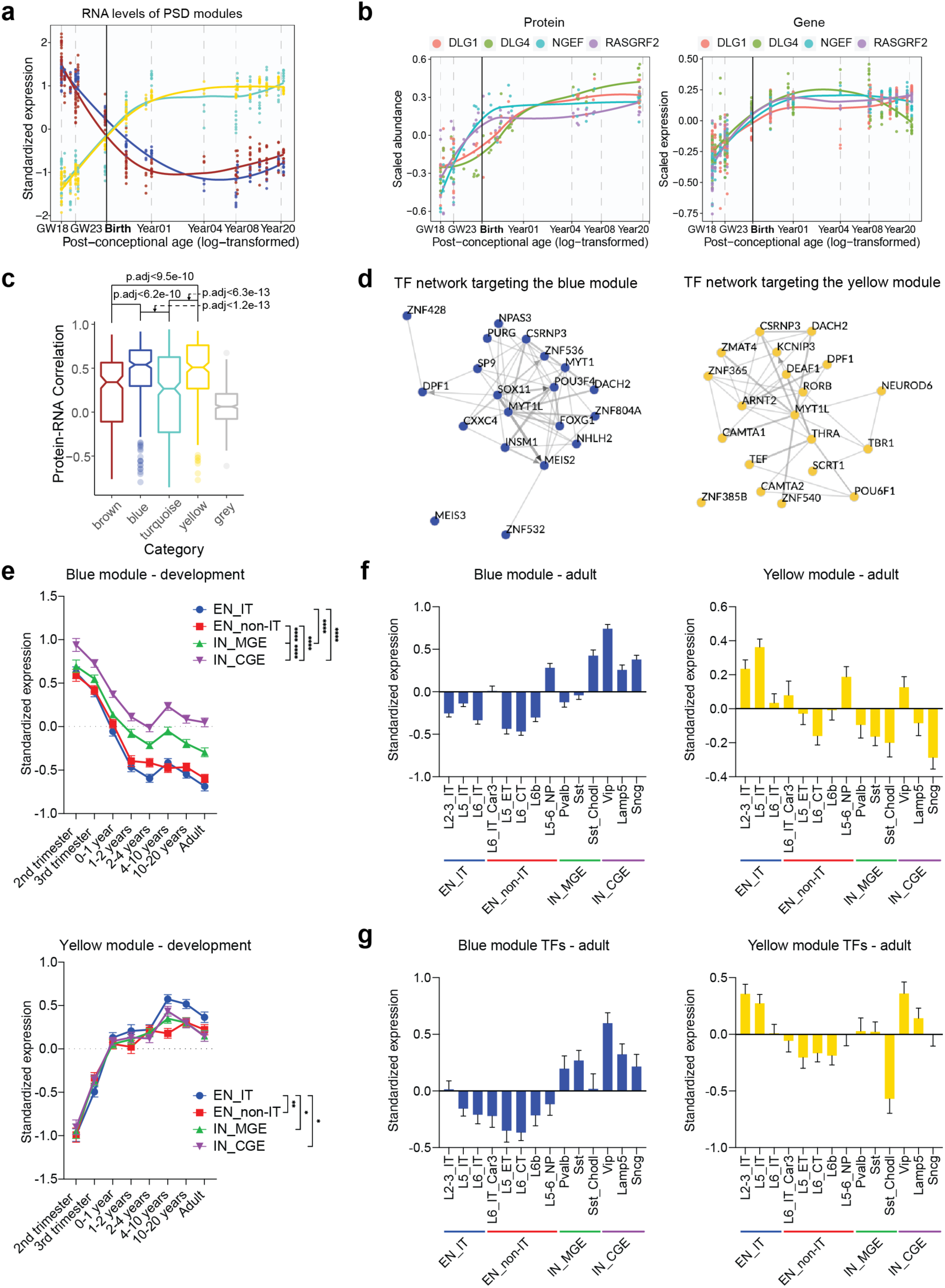
Transcription of PSD proteins and cell type specificity. **a**, Standardized median expression values of genes encoding proteins of the four PSD modules in the BrainSpan data. **b**, Scaled protein abundance and gene expression patterns of DLG1, DLG4, NGEF, and RASGRF2. **c**, Spearman correlation coefficients between protein abundance and gene expression of PSD proteins in each module. **d**, Transcription factor (TF) networks that regulate genes in the blue and yellow modules. **e**, Standardized expression values of genes in the blue and yellow modules in individual neuronal subtypes of the developing human neocortex. EN_IT: excitatory intratelencephalic neuron; EN_non-IT: excitatory non-intratelencephalic neuron; IN_MGE: inhibitory neuron derived from the medial ganglionic eminence; IN_CGE: inhibitory neuron derived from the caudal ganglionic eminence. **f**, Standardized expression values of genes in the blue and yellow modules in individual neuronal subtypes of the adult human neocortex. **g**, Standardized expression values of TFs predicted to regulate the blue and yellow modules in individual neuronal subtypes of the adult human neocortex.

The high RNA-protein concordance of proteins in the blue and yellow modules suggests that their PSD abundances are mainly regulated by transcription. Therefore, we focused on the blue and yellow modules to study the transcriptional regulatory mechanisms of PSD development. Transcription factor (TF) enrichment analysis by ChEA3 revealed core TF networks targeting the two modules (Fig. 3d and Supplementary Table 6). Some of the TFs identified in these networks, such as FOXG1, MEIS2, MYT1L, and RORB, are known to be critical regulators of neuronal differentiation and synapse development. To investigate transcription of the blue and yellow modules in a cell type-specific manner, we integrated our PSD proteomic data with single-cell RNA-sequencing data from the developing and adult human neocortex (Extended Data Fig. 12b,c, Supplementary Table 7) ^39,40^. We found that both excitatory neurons (EN) and inhibitory neurons (IN) had a reduction in blue module gene expression and an increase in yellow module gene expression during development. However, the developmental reduction of blue module genes in INs, particularly those derived from the caudal ganglionic eminence (IN_CGE), was slower than that of ENs (Fig. 3e). On the other hand, yellow module gene expression increased significantly faster in excitatory intratelencephalic neurons (EN_IT) (Fig. 3e). While there was more heterogeneity among individual neuronal subtypes in the adult brain, the overall trend remained consistent, with INs maintaining higher expression of genes encoding early-stage synaptic proteins compared to ENs (Fig. 3f). This difference in expression can be attributed to the differential expression of TFs targeting the two modules (Fig. 3g). Differences in transcription and abundance of PSD proteins may contribute to the distinct excitatory postsynaptic responses observed in these two subclasses of cortical neurons ^41^.

### Species differences in PSD development

Excitatory synapses and the PSD in humans, macaques, and mice are similar, yet they differ at both the morphological and molecular levels ^22,24,42^. In light of this, we sought to investigate the changes in PSD development that contribute to the differences across these three species. We collected macaque and mouse neocortical samples at five time points (Fig 4a, Supplementary Table 7). These time points roughly correspond to the developmental stages of our human samples^43^. Samples from macaques were collected from the PFC, as it was the predominant source of our human samples. Because mice do not have a granular PFC ^44^ and our findings from the human PFC can be generalized to other cortical areas, we collected whole mouse neocortex instead.

**Fig. 4.**
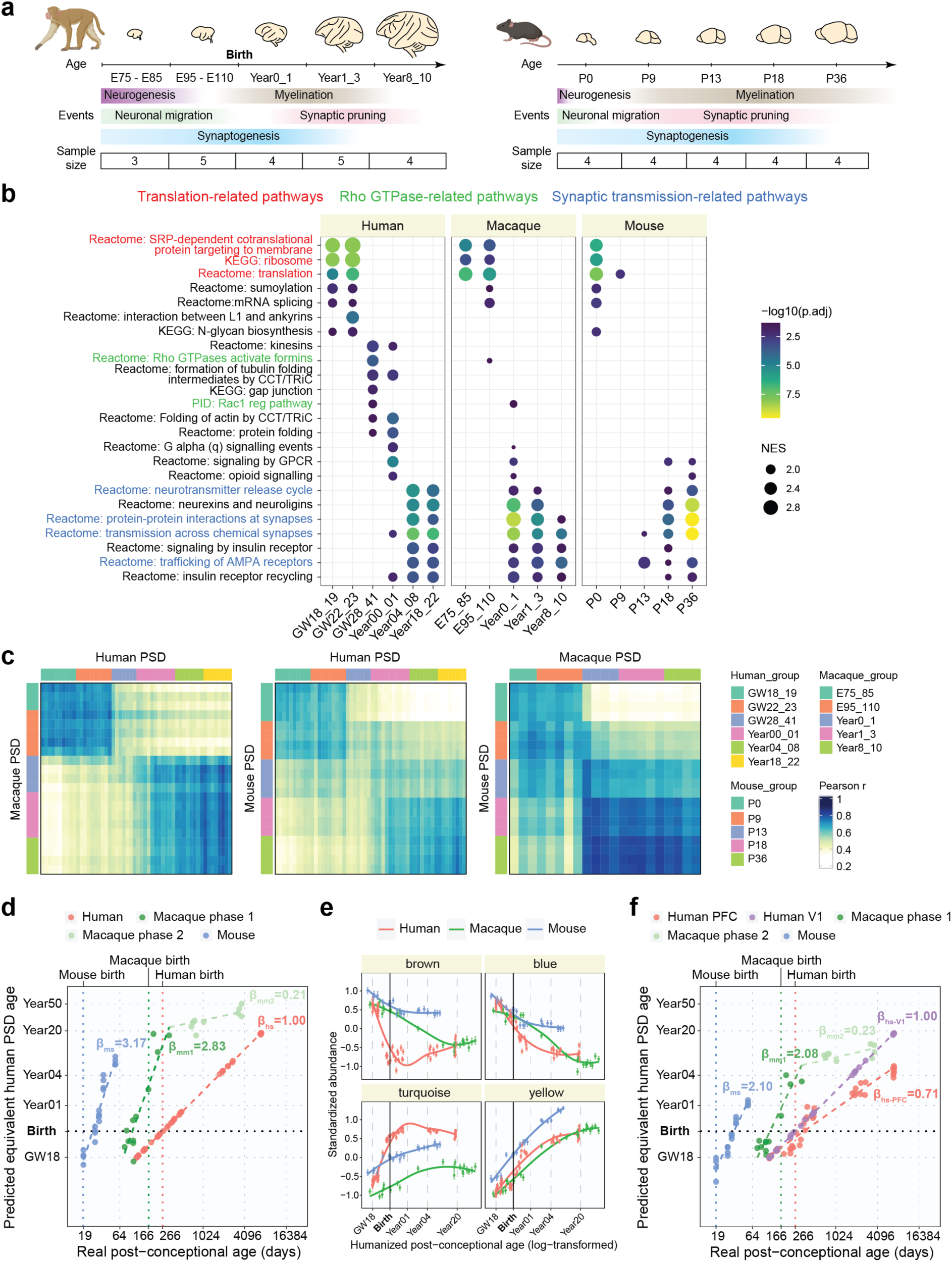
Comparison of PSD development across humans, macaques, and mice. **a**, Schematic illustrating the developmental stages of macaque and mouse samples. **b**, Gene set enrichment analysis for individual age groups across species. NES: normalized enrichment score. **c**, Similarity matrices representing pairwise Pearson correlations between human, macaque, and mouse samples. **d**, Predicted equivalent human PSD ages. β indicates the slope coefficients of the linear regression models in each species. **e**, Standardized abundance patterns of proteins in the four PSD modules across species. **f**, Predicted equivalent human PSD ages based on the human V1 dataset. β indicates the slope coefficients of the linear regression models in each region and species.

Proteomic analysis identified 1572 proteins in the developing macaque PSDs and 1572 proteins in mouse PSDs (Supplementary Tables 9,10), with some proteins being stage-specific (Extended Data Fig. 13a,b). Both PC analysis and hierarchical clustering showed that samples clustered by their age groups (Extended Data Fig. 13c–f). GSEA showed that, as in humans, translation-related pathways and synaptic transmission-related pathways were more active in early and late PSD development, respectively, in both macaques and mice (Fig. 4b). However, enrichment of pathways in the human perinatal phase, including Rho GTPase signaling and protein folding, was largely diminished in macaques and mice (Fig. 4b). To compare PSD samples from different species quantitatively, we performed cross-species similarity analysis. We calculated the Pearson correlation coefficients between PSD samples from different species and found that human samples in the second trimester and above four years of age correlated well with macaque and mouse samples at the corresponding stages (Pearson r > 0.6) (Fig. 4c). However, human samples between the third trimester and one year of age (the perinatal phase) showed relatively low correlations with all age groups in other species. We then sought to identify changes in PSD development that led to this difference.

Different species have distinct developmental timescales, making it hard to compare the abundance of PSD proteins directly. We, therefore, applied a regularized linear approach (see Methods) to unbiasedly predict the equivalent PSD ages of all three species based on their proteomic profiles (Fig. 4d, Supplementary Table 11). We found that multiplicative changes in real age were approximately linearly associated with multiplicative changes in the predicted equivalent human PSD age, except that macaque samples appeared to undergo two different stages separated by one year of age (Fig. 4d). We thus regressed the log-transformed humanized ages against the real log-transformed ages using a linear model (or a linear spline model for macaque samples) and obtained the slope coefficients as an estimator of PSD maturation rate normalized to the developmental timescale of individual species. This analysis revealed that PSD maturation was about three times slower in humans than in mice and macaques (< 1 year) (Fig. 4d).

Based on the equivalent PSD ages, we compared the abundance patterns of the human PSD modules in all three species. While the patterns of the blue and yellow modules were similar across all species, the brown and turquoise modules displayed species-specific differences (Fig. 4e). Specifically, the brown module was less abundant, and the turquoise module was more abundant in humans at the perinatal phase, likely causing the low correlation we observed in the similarity analysis at this developmental stage. Accordingly, module density and connectivity preservation analysis showed that the brown and turquoise modules were among the least preserved in macaques and mice (Extended Data Fig. 13g,h). Given the low RNA-protein concordance of these two modules (Fig. 3c), our results also highlight the role of post-transcriptional/translational regulation in shaping the distinctive features of human synapses.

Similar results were obtained when comparing the macaque and mouse datasets to the human V1 dataset (Fig. 4g, Extended Data Fig. 14). One minor but interesting difference is that the predicted PSD maturation rate in human V1 was about two times slower than in mice and macaques (< 1 year) and about 40% faster than in human PFC (Fig. 4f). This is consistent with previous findings that sensory cortical areas sensory cortex, such as V1, generally matures faster than association areas like PFC ^45,46^. In conclusion, human PSD matures at a slower rate in the neocortex, and the perinatal phase of its development is less represented in macaques and mice.

### Enhancement of RhoGEF signaling promotes neoteny of human synapses

The slower maturation rate of the human PSD could result from the increased abundance of turquoise module proteins and enrichment of Rho GTPase regulators at the perinatal phase. To test this hypothesis, we further investigated the increase of RhoGEF signaling in the human PSD. Indeed, most RhoGEF proteins in the turquoise module were greatly increased at the perinatal phase in humans and remained more abundant than those in other species thereafter in our proteomic data (Fig. 5a). This finding was further confirmed by Western blotting (Fig. 5b, Extended Data Fig. 15a–c) and immunostaining (Fig. 5c, Extended Data Fig. 16). Note that no significant changes were observed in the total homogenate over time (Extended Data Fig. 15a), consistent with our hypothesis that protein trafficking plays a crucial role in regulating the PSD abundance of turquoise module proteins. Postmortem accumulation could lead to an artificial increase in PSD proteins as has been reported for tubulins ^47^. To rule out the possibility that postmortem accumulation caused the observed increase in RhoGEF proteins, we compared adult PSDs prepared from postmortem samples (postmortem interval between 14 to 17 hours) with those from neurosurgical biopsy and found that RhoGEF levels were comparable (Extended Data Fig. 15d). Next, we tested whether the increase in RhoGEF proteins led to the activation of their known downstream pathways. A majority of RhoGEF proteins in the turquoise module target Rac1. We found that phosphorylation of PAK and CFL1, both downstream effectors of Rac1, increased in human synaptosomes along with RhoGEFs during synapse maturation (Extended Data Fig. 15e,f). However, no change was observed in mouse neurons (Extended Data Fig. 15e,f). Together, these results validated the enhancement of RhoGEF signaling in the human PSD.

**Fig. 5.**
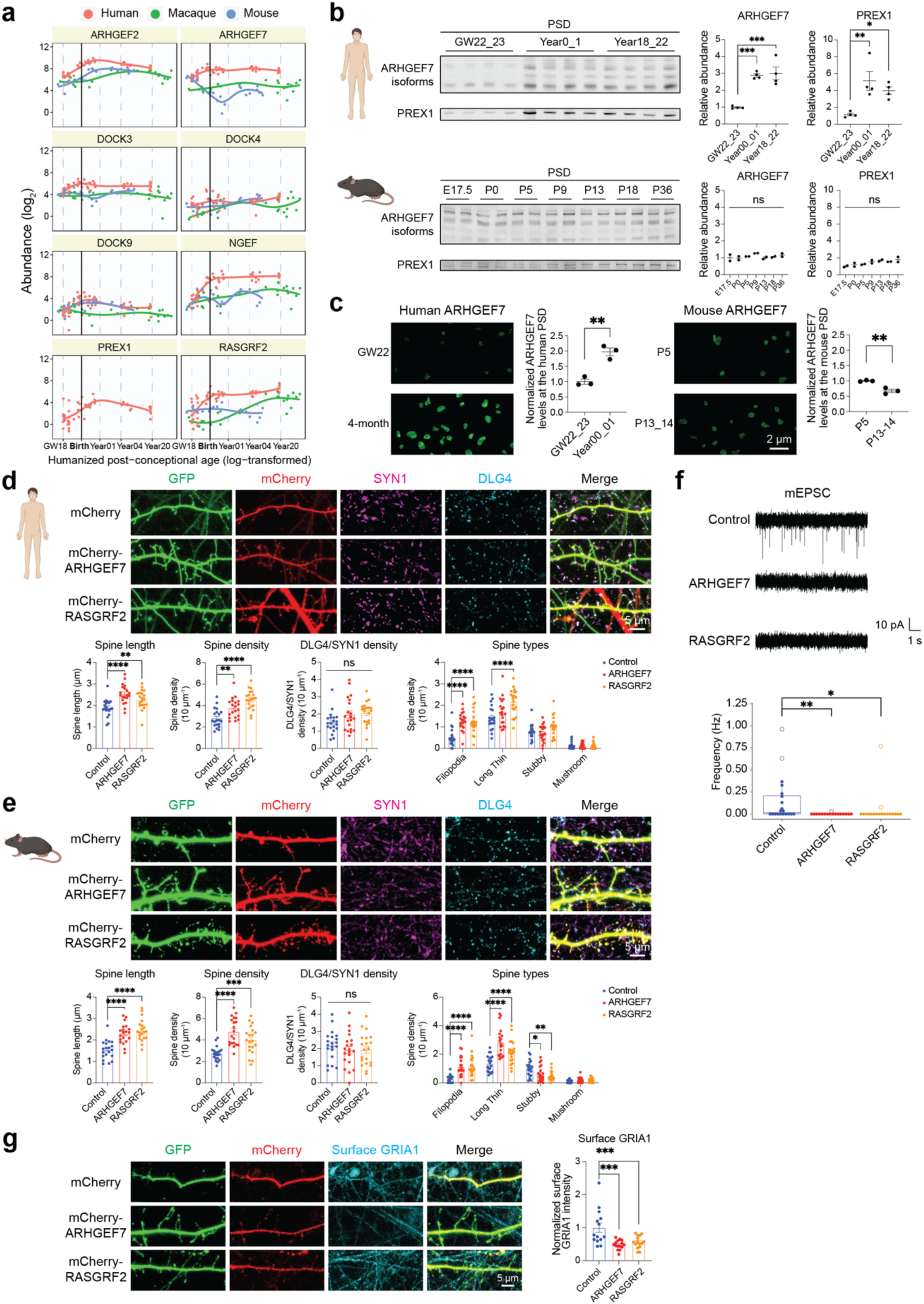
Increase in RhoGEF proteins promotes neoteny of human synapses. **a**, Abundance patterns of RhoGEFs in the turquoise module across species. **b**, Immunoblots and quantification of representative RhoGEFs in the PSD of the developing human (n = 4, 4, 4 samples) and mouse (n = 2, 2, 2, 2, 2, 2, 2 samples) PSD. *p < 0.05, **p < 0.01, ***p < 0.001; one-way ANOVA with Holm-Sidak’s multiple comparisons test. **c**, Immunofluorescent intensity of ARHGEF7 at DLG4 loci in the developing human and mouse neocortex (n = 3, 3, 3, 3 samples, scale bar: 2 μm). **p < 0.01; unpaired two-tailed *t* test. **d**, Immunostaining of dendrites from primary human cortical neurons cultured six weeks *in vitro* transfected with mEGFP-C1 and vectors expressing mCherry, mCherry-ARHGEF7, or mCherry-RASGRF2 (n = 20, 20, 20 neurons, scale bar: 5 μm). **p < 0.01, ****p < 0.0001; one-way ANOVA with Holm-Sidak’s multiple comparisons test. **e**, Immunostaining of dendrites from primary mouse cortical neurons cultured 8 days *in vitro* transfected with mEGFP-C1 and vectors expressing mCherry, mCherry-ARHGEF7, or mCherry-RASGRF2 (n = 20, 20, 20 neurons, scale bar: 5 μm). *p < 0.05, **p < 0.01, ***p < 0.001, ****p < 0.0001; one-way ANOVA with Holm-Sidak’s multiple comparisons test. **f**, Miniature excitatory postsynaptic current (mEPSC) recording of primary human cortical neurons cultured six weeks *in vitro* transfected with mEGFP-C1 and vectors expressing mCherry, mCherry-ARHGEF7, or mCherry-RASGRF2 (n = 20, 17, 18 neurons). *p < 0.05, **p.adj < 0.01; Kruskal–Wallis test with Dunn’s multiple comparisons test. **g**, Immunostaining against surface GRIA1 of dendrites from primary human cortical neurons cultured six weeks *in vitro* transfected with mEGFP-C1 and vectors expressing mCherry, mCherry-ARHGEF7, or mCherry-RASGRF2 (n = 14, 15, 15 neurons, scale bar: 5 μm). ***p < 0.001; one-way ANOVA with Holm-Sidak’s multiple comparisons test.

Next, we manipulated Rho GTPase regulators in neurons to understand their role in synapse maturation. We individually overexpressed two RhoGEF proteins from the turquoise module, ARHGEF7 and RASGRF2, in developing human cortical neurons (Extended Data Fig. 17). The density of synapses, quantified by DLG4 and SYN1 co-staining, was similar (Fig. 5d), indicating that synaptogenesis was not affected. However, analysis of the morphology of dendritic spines revealed that overexpressing either ARHGEF7 or RASGRF2 increased spine length and promoted the formation of more immature spine types (long thin and filopodia) (Fig. 5d). We also observed a significant increase in spine density in RhoGEF overexpressing neurons (Fig. 5d). Similar results were observed in mouse cortical neurons (Fig. 5e). Conversely, individually knocking down two RhoGAP proteins from the blue module, ARHGAP23 and SRGAP1, partially phenocopied RhoGEF overexpression in increasing spine length and immaturity (Extended Data Fig. 18, Supplementary Table 12). To test whether these morphological changes translate into functional consequences, we recorded miniature excitatory postsynaptic currents (mEPSCs) in human neurons with and without RhoGEF overexpression. We found that mEPSC frequency was significantly decreased in neurons overexpressing either ARHGEF7 or RASGRF2 (Fig. 5f). Moreover, the surface level of AMPA receptor GRIA1 was reduced by ARHGEF7 or RASGRF2 overexpression (Fig. 5g, Extended Data Fig. 19). These data suggest that overexpression of specific RhoGEF proteins increases filopodial silent synapses and inhibits their functional maturation. Altogether, our results suggest that the human-specific increase in selective RhoGEF proteins delays maturation and promotes neoteny of human synapses.

### PSD modules in human cognition and brain disorders

We next investigated whether genetic variants associated with human cognition converge onto the human PSD modules, identifying that the turquoise module was enriched for GWAS signals of processing speed and fluid intelligence (the UK Biobank) (Fig. 6a). The turquoise module has its peak expression shortly after birth, at which time infants perceive a wealth of external stimuli. Thus, we posited that proteins in this module were important for activity-dependent synaptic remodeling. Indeed, the turquoise module was highly enriched for activity-dependent proteins in neurons (Odds ratio > 3; Fig. 6b, Supplementary Table 13) ^48^, including TRIM3 (an activity-dependent ubiquitin ligase for PSD scaffolding proteins) ^49^, GRIPAP1 (a recycling endosome regulator critical for synaptic plasticity) ^50^, and NGEF (a RhoGEF protein) (Extended Data Fig. 20). Combined with the fact that the turquoise module is more abundant in the human PSD and that RhoGEF proteins in this module promote synaptic neoteny, these results highlight the possible significance of this module in the evolutionary enhancement of human cognitive function.

**Fig. 6.**
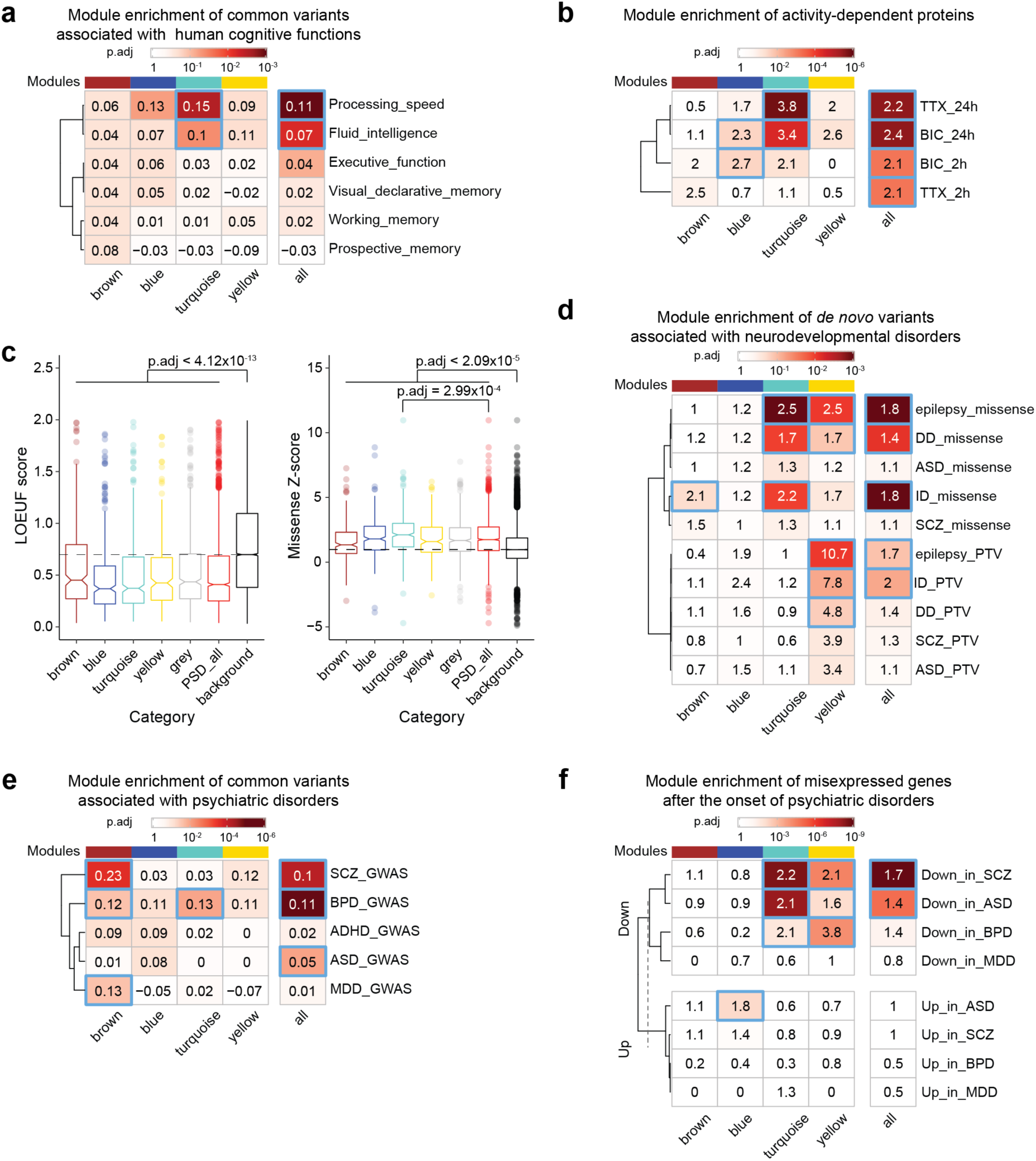
Association of human PSD modules with cognitive functions and brain disorders. **a**, Enrichment of common variants associated with human cognitive functions in PSD modules. The numbers indicate the MAGMA linear regression coefficient β. The blue borders denote p.adj < 0.05; MAGMA analysis on GWAS summary statistics. **b**, Enrichment of neuronal activity-dependent proteins in PSD modules. The numbers indicate the odds ratio. The blue borders denote p.adj < 0.05; hypergeometric test. **c**, Distribution of gnomAD LOEUF scores and missense Z-scores of genes in each category. Kruskal–Wallis test. **d**, Enrichment of de novo variants associated with neurodevelopmental disorders in PSD modules. The numbers indicate the odds ratio. The blue borders denote p.adj < 0.05; hypergeometric test. **e**, Enrichment of common variants associated with psychiatric disorders in PSD modules. The numbers indicate the MAGMA linear regression coefficient β. The blue borders denote p.adj < 0.05; MAGMA analysis on GWAS summary statistics. **f**, Enrichment of misexpressed genes after the onset of psychiatric disorders in PSD modules. The numbers indicate the odds ratio. The blue borders denote p.adj < 0.05; hypergeometric test.

Synaptic dysfunction contributes to both neurodevelopmental and psychiatric disorders, often caused by *de novo* and common variants, respectively. Regarding *de novo* variants, genes encoding PSD proteins were more intolerant of protein-truncating variants (PTVs) (lower LOEUF scores) and missense variants (higher missense Z-scores) compared with all genes expressed in the neocortex (Fig. 6c), suggesting that mutations in these genes are more likely to cause human diseases. Remarkably, the turquoise module was particularly intolerant of missense variants (Fig. 6c). No difference was observed for synonymous mutations (Extended Data Fig. 20b). Accordingly, we found that genes encoding PSD proteins were enriched for *de novo* nonsynonymous variants associated with neurodevelopmental disorders including epilepsy, developmental delay (DD), and intellectual disability (ID) (denovo-db) (Fig. 6d, Extended Data Fig. 20c,d, Supplementary Table 13). In particular, the turquoise module had excessive missense variants, whereas the yellow module was enriched for both missense variants and PTVs. Turquoise module genes with disease-associated missense variants included genes encoding ion channels such as *KCNQ2* and *SCN2A* and molecular motors such as *DYNC1H1* and *KIF1A*. Some of these missense variants may be gain-of-function or dominant-negative mutations with different pathogenic effects than PTVs ^51,52^. In contrast, many yellow module genes with PTVs were genes encoding enzymes that regulate PSD organization and postsynaptic receptor trafficking, such as *SYNGAP1*, *IQSEC2*, and *CDKL5*. Therefore, although mutations in both modules contribute to neurodevelopmental disorders, the different patterns of module abundance and variant type enrichment suggest that they do so by different mechanisms that target distinct stages of synapse maturation.

For psychiatric disorders, the brown module was enriched for GWAS signals of diseases that generally manifest in young adulthood, including schizophrenia (SCZ), bipolar disorder (BPD), and major depressive disorder (MDD) (The Psychiatric Genomics Consortium) (Fig. 6e, Extended Data Fig. 21a, Supplementary Table 13). Proteins in the brown module had peak abundance at midgestation, indicating an early etiology involving synapse development for these adolescence/adult-onset disorders. However, after the onset of these psychiatric disorders, genes encoding PSD proteins in the late modules (turquoise and yellow) were downregulated compared with controls (Fig. 6f, Extended Data Fig. 21b,c, Supplementary Table 13). This is likely the consequence of a downstream cascade of biological events following earlier-acting genetic risk factors that disrupt synapse development.

## Discussion

Although synapses and the PSD are known to undergo profound remodeling in brain development to enable the formation and reorganization of brain networks ^10,12,53^, we have had limited knowledge of the molecular changes that occur during this remodeling. In this study, we generated a cross-species proteomic map of synapse development and revealed the temporal dynamics of >1,000 PSD proteins. We demonstrate that the human PSD undergoes three major phases of maturation. By relating the abundance of PSD proteins to each other, we further uncovered individual protein modules and networks that exert stage-, cell type-, and species-specific functions. Furthermore, we found that the PSD develops about two to three times slower in human neocortex than in other species and that the increased abundance of RhoGEF proteins, as expressed in the turquoise module, contributes to this difference. The turquoise module is also associated with synaptic plasticity, human cognitive function, and mental disorders. Together, these data provide a blueprint for studying the molecular and evolutionary mechanisms of synapse maturation in humans.

Synapse development is regulated at both RNA and protein levels ^54^. By integrating PSD proteomic data with bulk RNA-sequencing data, we found that different PSD modules exhibit different RNA-protein concordance, suggesting that they are differentially regulated by post-transcriptional/translational mechanisms. Focusing on the modules with high RNA-protein concordance, we inferred neuronal subtype-specific PSD signatures from single-cell RNA-sequencing data. Our analysis revealed major differences in the PSD between excitatory and inhibitory neuronal subtypes in the neocortex. This is consistent with previous studies showing that the composition of the PSD is diverse among neuronal subtypes ^55–57^. Moreover, INs have higher levels of blue module genes, including those encoding RhoGAPs which suppress dendritic spine formation. This could contribute to the differences in spine densities observed between ENs and INs ^58^. One limitation of this inference is that single-cell RNA-sequencing does not include RNAs in dendrites that could contribute to the PSD through local translation. Although somatic and dendritic RNAs are significantly correlated ^59^, future studies to determine the proteomic profiles of neuronal subtype-specific PSDs will help expand these findings.

Previous studies identified a critical role of a RhoGAP protein, SRGAP2, in the human-specific developmental delay in synapse maturation and increase in synaptic density ^60–62^. It has been shown that the Rac1-GAP activity of the ancestral protein SRGAP2A limits the spine neck length and density in neocortical neurons. Human-specific partial duplications of SRGAP2 inhibited the function of SRGAP2A, resulting in longer spine necks and higher spine density in humans. Our study found that multiple RhoGEF proteins have increased abundance in the human PSD starting at the perinatal stages. Enhancement of RhoGEF signaling in neurons not only increased spine length and density but also delayed functional maturation of synapses. Given the antagonistic roles of RhoGAP and RhoGEF proteins in activating Rho GTPases, our results are consistent with the previous findings and suggest that increased synaptic Rho GTPase activity contributes to the neoteny of human synapses. Moreover, RhoGEF proteins are enriched in the turquoise module associated with human cognitive function. Thus, our analysis provides molecular evidence that links synaptic neoteny to the evolution of human cognition.

Early synaptic connections before the third trimester are often transient stepping-stones toward functional synaptic circuits in mature brains ^63^. Surprisingly, genetic variations of PSD proteins specifically abundant at midgestation are associated with adolescent-onset psychiatric disorders. Given the dysregulation of late-stage synaptic proteins after the onset of these disorders, our findings highlight the importance of early synaptic connections for shaping neuronal wiring and higher-order brain functions of the mature brain.

There are several limitations to this study. Isolation of PSDs by subcellular fractionation can include contaminants from other cellular compartments or associated structures ^64^. We have therefore applied multistep orthogonal data filtering to minimize the effect of contamination. Additional independent validation, such as proximity proteomics or immunogold labeling, will further determine if newly identified proteins are *bona fide* PSD components. Moreover, alternative splicing and the isoforms they produce play a key role in regulating synapse development ^65,66^. However, our proteomic analysis does not include quantifications at the isoform level due to technical limitations. Furthermore, PSDs are heterogenous and likely different between individual synapses, yet our data represent the averaged proteomic profiles of PSDs at the bulk tissue level. Lastly, the enrichment of activity-dependent proteins in the turquoise module was based on experimental data from rat neurons but not human neurons ^48^. With the development of novel methods, future studies determining developmental and activity-dependent changes of the synaptic proteome at the isoform level across different brain regions, cell types, and species will provide further insight into the mechanisms of brain development, evolution, and disease.

## Methods

### Brain tissue samples

Acquisition of all second-trimester primary human tissue samples (Supplementary Table 1a) was approved by the UCSF Human Gamete, Embryo and Stem Cell Research Committee (study number 10-05113). All experiments were performed in accordance with protocol guidelines. Informed consent was obtained before sample collection and use for this study. After tissue acquisition, the cortical plate and subplate were dissected and frozen at −80 °C.

Thirty-four de-identified snap-frozen post-mortem prefrontal cortex (PFC) tissue samples without known neurological disorders were obtained from the University of Maryland Brain and Tissue Bank through NIH NeuroBioBank and the Pediatric Brain Tissue Bank at UCSF (Supplementary Table 1a).

Twenty-two de-identified snap-frozen post-mortem primary visual cortex (V1) tissue samples without known neurological disorders were obtained from the University of Maryland Brain and Tissue Bank through NIH NeuroBioBank (Supplementary Table 1b).

Three adult cortical samples were obtained from neurosurgical operations (Supplementary Table 1c). In two cases, the reason for surgery was tumor resection and in one, focal cortical dysplasia type 2b treatment. In all cases, the tissue samples were collected from peripheral regions removed together with the affected regions. The samples were snap-frozen in liquid nitrogen and stored at −80 °C before further processing.

Macaque samples were obtained from Oregon National Primate Research Center, Southwest National Primate Research Center, Wisconsin National Primate Research Center, Dr. Alex Pollen at UCSF, and Dr. Nenad Sestan at Yale University (Supplementary Table 8a).

Mouse experiments were approved by UCSF Institutional Animal Care and Use Committee (IACUC) and performed in accordance with relevant institutional guidelines. Neocortices from C57BL/6J mice were dissected and frozen at −80 °C (Supplementary Table 8b).

### Subcellular fractionation and post-synaptic density (PSD) isolation

Subcellular fractionation and PSD isolation were done as previously described ^29^. Briefly, about 300 to 2000 mg of human brain tissue (dependent on the age group) was thawed on ice and cut into small pieces. It was then homogenized using a 10 mL tissue grinder in 3 mL homogenization buffer (0.32 M sucrose, 2 mM EDTA, and 4 mM HEPES, pH 7.4) with freshly added protease and phosphatase inhibitors (Roche). Postnuclear supernatants were obtained via 1,000 × g spin. The supernatant (S1) was spun at 10,000 × g for 15 min, and the pelleted membranes were resuspended in 10 mL homogenization buffer and spun again to obtain P2’. This pellet was lysed by 10 mL hypoosmotic shock in 4 mM HEPES (pH7.4) buffer and incubated for 30 min at 4 °C. This was followed by a spin at 25,000 × g for 20 min. The pellet (P3) was layered onto a 0.8/1.0/1.2 M discontinuous sucrose gradient and spun at 150,000 × g for 2 h. Synaptic plasma membranes (SPM) were recovered from the 1.0/1.2 M interface and pelleted. These membranes were extracted with 3 mL 0.5% Triton X-100 for 15 min and pelleted with a 32,000 × g spin for 20 min (PSD I). For mass spectrometry (MS) and Western blot analysis, fractions obtained by subcellular fractionation were lysed by sonication. Protein concentrations were determined using the BCA assay (Pierce).

Macaque and mouse PSD samples were prepared in the same way.

### Western blot

LDS sample buffer (Invitrogen) with a reducing agent was added to each protein lysate, followed by a 10 min incubation at 95 °C. Samples were spun down and electrophoresed on a 4–12% Bis-Tris gel and transferred to a nitrocellulose membrane. Total protein was quantified by Revert Total Protein Stains (LI-COR). The membrane was then blocked for 1 h with Intercept (TBS) Blocking Buffer (LI-COR) before primary antibody incubation overnight. After secondary antibody incubation, the membrane was imaged using Odyssey Classic or Odyssey CLx (LI-COR).

The following antibodies were used: GRIN2B (Abcam, ab93610, 1:1,000), DLG2 (NeuroMab, 75-284, 1:1,000), GRIA1 (Millipore, AB1504, 1:1,000), DLG4 (NeuroMab, 75-028, 1:1,000), SYP (Sigma, S5768, 1:1,000), GAPDH (Cell Signaling Technology, 2118S, 1:1,000), ARHGEF7 (Sigma, 07-1450-I, 1:1,000), PREX1 (Sigma, HPA001927-100UL, 1:1000), β-Tubulin (Cell Signaling Technology, 2146S, 1:1,000), Phospho-PAK (Cell Signaling Technology, 2606S, 1:1,000), PAK (Cell Signaling Technology, 2604S, 1:1,000), Phospho-CFL (Cell Signaling Technology, 3311S, 1:1000), CFL (Cell Signaling Technology, 5175S, 1:1000).

### Transmission electron microscopy

Human PSD I pellets prepared from gestational week (GW) 23 samples were fixed with 3% glutaraldehyde (Electron Microscopy Sciences, EMS) for 10 min at 4 °C, washed three times with 0.1M phosphate buffer (PB), and post-fixed with 2% osmium tetroxide (EMS) in 0.1M PB for 1 h at room temperature. Dehydration in graded series of ethanol (30, 50, and 70%) was followed by 2% uranyl acetate (EMS) incubation for 2.5 h. Samples were rinsed with 70%, 96%, and 100% ethanol, washed two times with propylene oxide (EMS), embedded in Durcupan resin (Sigma-Aldrich), and allowed to polymerize at 69 °C for 72 h. Ultrathin sections (60–70 nm) were sectioned with a diamond knife (DiATOME) in a UC7 ultramicrotome (Leica), stained with lead citrate, and examined under a transmission electron microscope (Tecnai Spirit G2, FEI, Netherlands) using a digital camera (Morada, Soft Imaging System, Olympus, Japan).

### Sample quality control, digestion, and LC-MS/MS analysis

The integrity of sample synaptic proteomes was checked before MS analysis using the HUman Synapse Proteome Integrity Ratio (HUSPIR) ^30^. PSD samples with HUSPIR > 2 were deemed good, and those with 1 < HUSPIR < 2 were deemed fair. Samples with HUSPIR ≤ 1 were excluded from further analysis.

Five micrograms of isolated PSD samples were electrophoresed on a 4–12% Bis-Tris gel for 20 min. Proteins were visualized by Bio-Safe Coomassie stain (BIO-RAD) and excised from the gel. Individual gel piece was subjected to in-gel digestion using trypsin (Promega). The resulting dried peptides were analyzed in technical duplicate on a ThermoFisher Orbitrap Fusion Lumos Tribrid mass spectrometry system equipped with an EASY-nLC 1200 ultrahigh-pressure liquid chromatography system interfaced via a Nanospray Flex nanoelectrospray source. Samples were injected into a C18 reverse phase column (25 cm x 75 µm packed with ReprosilPur C18 AQ 1.9 µm particles). Peptides were separated by an organic gradient from 5 to 30% ACN in 0.02% heptafluorobutyric acid over 180 min at a flow rate of 300 nL/min for the phosphorylated peptides or unmodified peptides for global abundance. Spectra were continuously acquired in a data-dependent manner throughout the gradient, acquiring a full scan in the Orbitrap (at 120,000 resolution with an AGC target of 400,000 and a maximum injection time of 50 ms) followed by as many MS/MS scans as could be acquired on the most abundant ions in 3 s in the dual linear ion trap (rapid scan type with an intensity threshold of 5000, HCD collision energy of 32%, AGC target of 10,000, maximum injection time of 30 ms, and isolation width of 0.7 m/z). Single and unassigned charge states were rejected. Dynamic exclusion was enabled with a repeat count of 2, an exclusion duration of 20 s, and an exclusion mass width of ±10 ppm.

### Protein identification, quantification, and potential contaminants removal

Protein identification and quantification were done using MaxQuant v1.6.11.0 ^67^. Spectra from the human, macaque, and mouse raw files were matched to the reference proteomes from UniProt (Homo sapiens UP000005640_9606, Macaca mulatta UP000006718_9544, and Mus musculus UP000000589_10090 respectively). Default settings of MaxQuant with FDR = 0.01 were used except that “Match between runs” was enabled to improve proteome coverage with an alignment window of 20 min and a match time window of 0.7 min. The iBAQ value for each protein group was calculated ^68^.

Potential external contaminants including keratins and proteins known to be localized at mitochondria ^69^, a principal contaminant in PSD preparation ^70^, were excluded. Moreover, because some identified proteins could be non-PSD proteins artificially bound to the PSD in the post-mortem condition ^47^, we curated a list of proteins that have been identified in non-post-mortem brain tissues by combining our data and data from Bayés et al., 2011 ^9^. This list was used to filter the identified PSD proteins so that those only present in post-mortem brain samples were excluded. After contaminant filtration, the remaining proteins were considered present in an age group if more than half of the samples in that group had the protein identified by MS/MS.

### Data normalization, imputation, and integrity effect correction

Our initial analysis focused on the human PFC dataset consisting of the 20 cortical samples without area information in the second trimester and 34 PFC samples from the third trimester to young adulthood. After filtering out potential contaminants, the iBAQ values of the remaining proteins were used to calculate the normalized molar intensity riBAQ ^71^:

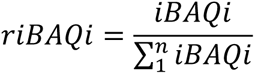

The riBAQ values were multiplied by a scale factor of 10^6^ and log2-transformed to obtain the abundance values for each protein. After log-transformation, 1765 human PSD proteins missing in less than three samples in any age group (either by MS/MS or by matching) were included for further analysis. Next, abundance values were normalized by variance stabilizing transformation, and missing values were imputed using the “MinProb” method (q = 0.01), both of which were implemented using the R package DEP ^72^.

Linear models combined with empirical Bayes methods were used for the differential abundance analysis of the human samples. We accounted for the fact that synaptic proteome integrity could affect the abundance values. The following model was fit:

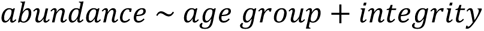

where *age group* is one of the six sample age groups, and *integrity* is the HUSPIR category (good or fair). This was done using the R package limma ^73^. After the model was fitted, a pairwise comparison was made to identify proteins with differential abundance between two age groups. Proteins with a log2 fold change of at least one and an adjusted p-value ^74^ of the moderated t-test less than 0.05 were selected as proteins with differential abundance. The effect of integrity was removed using the removeBatchEffect function in limma. The final corrected abundance matrix can be found in Supplementary Table 2a.

A similar differential abundance analysis was done with the macaque and mouse samples, except that the integrity covariate was removed in the linear model. The final abundance matrix can be found in Supplementary Table 9a and Supplementary Table 10a.

A similar differential abundance analysis was done with the human V1 dataset consisting of the same 20 cortical samples without area information in the second trimester as the PFC dataset and 22 V1 samples from the third trimester to young adulthood. Variance stabilizing transformation was done using the same model from the PFC dataset. The final corrected abundance matrix after applying the removeBatchEffect function can be found in Supplementary Table 5a.

### Dimensionality reduction and clustering

Dimensionality reduction was done by the principal component analysis (PCA). The first two principal components were used for the PCA plots. Heatmaps were generated using the R package ComplexHeatmap ^75^. Samples and proteins were clustered based on the Spearman correlation distance.

### Variance partitioning

Variance partitioning was done using the R package variancePartition ^76^. Age group, processing batch, PSD quality, and sex of the sample donor were included in the formula.

### Gene set enrichment analysis using MSigDB gene sets

Age specificity for each protein was calculated by comparing samples within an age group to all other samples outside the age group using linear models combined with empirical Bayes methods implemented in limma. The resulting moderated t statistics of each protein were ranked and used as input for gene set enrichment analysis (GSEA) ^77^ using the R package clusterProfiler ^78^. GSEA was carried against the MSigDB C2 canonical pathways, which contain curated gene sets representing different molecular pathways ^79^. Only pathway sets with gene numbers between 10 and 500 were used for the analysis.

### Immunohistochemistry and confocal imaging

Prenatal human tissue samples and mouse tissue samples were fixed in 4% paraformaldehyde in PBS at 4 °C overnight. The samples were cryoprotected in 15% and 30% sucrose in PBS and frozen in OCT. Samples were sectioned at a thickness of 15 µm, air-dried, and rehydrated in PBS. Postnatal unfixed frozen tissue samples were sectioned at a thickness of 15 µm, air-dried, fixed in 4% paraformaldehyde in PBS for 10 min, and washed three times with PBS. Antigen retrieval was done using citrate-based Antigen Unmasking Solution (Vector Laboratory) at 95 °C for 10 min. The slides were then washed in PBS and blocked in PBS-based blocking buffer containing 10% donkey serum, 0.2% gelatin, and 0.1% Triton X-100 at room temperature for 1 h. After blocking, slides were incubated with primary antibodies in the blocking buffer at 4 °C overnight. The slides were washed in PBS three times and incubated with secondary antibodies in the blocking buffer at room temperature for 2 h. The slides were then washed in PBS twice, counterstained with DAPI, and washed in PBS once more. Slides were mounted with coverslips with ProLong Gold (Invitrogen). Confocal images were acquired with a Leica TCS SP8 using a 63× oil immersion objective. Acquired images were processed using Imaris (Oxford Instruments) and Fiji ^80^. The following antibodies were used: DLG4 (NeuroMab, 75-028, 1:250), DLG4 (Synaptic Systems, N3702-At488-L, 1:250), RPS6 (Cell Signaling Technology, 2217S, 1:200), CTNNB1 (Cell Signaling Technology, 8480S, 1:100), GDI1 (Proteintech, 10249-1-AP, 1:100), CFL1 (Cell Signaling Technology, 5175S, 1:250), DBN1 (Abcam, ab11068, 1:200), DNBL (proteintech, 13015-1-AP, 1:100), ARHGEF7 (Sigma, 07-1450-I, 1:200).

Colocalization analysis was done using Imaris. Protein puncta were identified based on signal intensity and local contrast. Puncta of two different proteins within 0.5 μm were considered co-localized. DBN1, DBNL, and ARHGEF7 abundances at the PSD were quantified using Imaris. The DLG4 puncta were identified based on signal intensity and local contrast. These DLG4 puncta, deemed as the PSD loci, were used to create a mask channel. Intensities of DBN1, DBNL, or ARHGEF7 were then quantified within the mask channel.

### Weighted gene co-expression network analysis (WGCNA)

WGCNA was done using the R package WGCNA ^34^.

For the human samples, the blockwiseModules function (power = 20, corType = “pearson”, networkType = “signed”, deepSplit = 2, minModuleSize = 45, reassignThreshold = 1e-6, mergeCutHeight = 0.15, minKMEtoStay = 0.3, numericLabels = F, pamRespectsDendro = F) was used to build a signed weighted correlation network. In total, four modules plus the grey module for unassigned proteins were identified. Module memberships and module eigengene values of the samples are available in Supplementary Table 3.

### Protein-protein interaction (PPI) enrichment analysis by Monte Carlo permutation tests

The PPI network consisting of interactions between human proteins and those between mouse proteins was downloaded from the BioGRID database (https://thebiogrid.org/) ^81^. We used the PPI network consisting of all the module genes in the human PSD as the background PPI network. For each module, we constructed the module protein interaction network by extracting the interactions connecting all the proteins in the module and calculated the observed number of interactions. We then randomly sampled the same number of proteins with similar degree distribution from the background PPI network and calculated the number of interactions in the random network. We repeated this randomization process 100,000 times and calculated the p-value as the fraction of the random numbers of interactions that are greater than the observed number of interactions.

### Statistical overrepresentation test using MSigDB gene sets

The one-sided hypergeometric test implemented in clusterProfiler was used to identify overrepresented pathways in each protein module. The MSigDB C2 canonical pathways were used as input gene sets. The union of genes encoding PSD proteins and those expressed in the human neocortex curated using the BrainSpan RNA-seq data was used as the background. To define genes expressed in the human neocortex, we downloaded the developing human brain RNA-Seq data from the BrainSpan database (http://www.brainspan.org). The neocortical samples of the BrainSpan data that have RNA integrity numbers ≥ 8 were temporally divided into prenatal and postnatal stages. Genes with RPKM ≥ 1 in at least half of the neocortical samples at either stage were defined as genes expressed in the human neocortex. Only pathway sets with gene numbers between 10 and 500 were used for the analysis.

### SynGO enrichment analysis

SynGO (release 1.1) enrichment analysis was done using the online tool at https://syngoportal.org/35. We used the same background gene list as the above statistical overrepresentation test.

### PPI-co-abundance network analysis and visualization

The pairwise topological overlap similarity scores of all human PSD protein pairs were calculated by WGCNA. Protein pairs with topological overlap similarity scores in the top 10% were considered co-expressed. Protein pairs that are co-expressed and interacting with each other in the BioGRID database were deemed connected in the PPI-co-abundance network. Subnetworks were generated within each module. All network plots were constructed using Cytoscape 3.8.2 ^82^. The shortest path lengths between proteins were calculated using igraph ^83^. The average length within pathway proteins was compared to that between the pathway proteins and non-pathway proteins in the network by the one-sided Wilcoxon rank-sum test.

### Protein domain analysis

Protein domain information of all PSD proteins was downloaded from the SMART database (https://smart.embl.de/) ^84^ and summarized in Supplementary Table 4a. Domains present in more than six PSD proteins were included in Fig. 2d and Extended Data Fig. 9. Domains were clustered based on the Pearson correlation distance in Fig. 2d and Jaccard distance in Extended Data Fig. 9.

### RhoGAP and RhoGEF specificity

The target specificity of individual RhoGAPs and RhoGEFs was hand-annotated based on a literature review. The annotation results were summarized in Supplementary Table 4b,c.

### Comparison between PSD proteomic data and bulk transcriptomic data

The BrainSpan human brain bulk RNA-seq data were downloaded from the PsychENCODE website (http://evolution.psychencode.org/) ^36^. Neocortical samples older than GW16 were included for analysis. Transcripts were filtered to only include those encoding PSD proteins and normalized to obtain the TPM values. Log2(TPM+1) values were used for downstream analysis.

Module preservation analysis was done using the modulePreservation function in WGCNA ^38^. We performed the permutation 500 times to obtain the Z_summary_ statistics and composite module preservation statistic medianRank.

To estimate the correlation at the individual PSD protein level between the proteomic data and transcriptomic data, we first imputed the transcriptomic profiles of human cortical samples used to generate our proteomic data. To do so, we trained a generalized additive model by regressing the transcriptome of human cortical samples against their ages using the R package mgcv 1.8-40^85^. We then predicted the corresponding transcriptomic profiles of our PSD samples based on their ages and calculated the Spearman correlation coefficients between RNA and protein of individual PSD proteins (Supplementary Table 6). The Kruskal-Wallis rank sum test with *post hoc* Dunn’s test was used to compare differences in correlation coefficients between PSD modules.

### ChEA3 transcription factor enrichment analysis

ChEA3 analysis was done using the online tool at https://maayanlab.cloud/chea3/ ^86^. Results were listed in Supplementary Table 6e.

### Developing human brain single-nucleus RNA sequencing (snRNA-seq) data analysis

SnRNA-seq data from the developing human neocortex ^87^ were downloaded from the UCSC Cell Browser ^88^. UMAP coordinates from the original authors were used. The identity of specific lineages and cell types was reannotated based on the expression of known marker genes (Extended Data Fig. 12). Only EN_IT, EN_non-IT, IN_MGE, and IN_CGE were included for further analysis. Pseudobulk samples were constructed by aggregating the raw counts of all the cells within the same cell type. Counts for genes encoding PSD proteins were extracted from the peudobulk data, normalized for sequencing depth, and log-transformed to obtain the log2(CPM+ 1) values (Supplementary Table 7a). Gene expression levels were standardized by z-transformation across pseudobulk samples and summarized in Fig. 3e.

### Adult human brain single-cell RNA sequencing (scRNA-seq) data analysis

scRNA-seq data from the adult human motor cortex ^40^ were downloaded from the UCSC Cell Browser ^88^. Only neocortical neurons were included for further analysis. Pseudobulk samples were constructed by aggregating the raw counts of all the cells within the same neuronal subtype annotated by the original authors. Counts for genes encoding PSD proteins were extracted from the peudobulk data, normalized for sequencing depth, and log-transformed to obtain the log2(CPM+ 1) values (Supplementary Table 7b). Counts for transcription factors regulating PSD modules were extracted from the original pseudobulk data without re-normalizing for sequencing depth (Supplementary Table 7c,d). Gene expression levels were standardized by z-transformation across pseudobulk samples and summarized in Fig. 3f,g.

### Correlation analysis between species and regions

The orthologs of genes encoding human PSD proteins were obtained from Ensembl. Only PSD proteins with one-on-one orthologs present in the PSD of all three species (854 proteins) were included for further analysis. The riBAQ values of the filtered data were re-normalized to obtain updated riBAQ values comparable between all species and regions (Supplementary Table 11a). Pairwise correlations between samples were obtained by calculating the Pearson correlation coefficients. The detailed results can be found in Supplementary Table 11.

### Prediction of equivalent human PSD ages and comparison of PSD maturation rates

To predict equivalent human PSD ages, we log-transformed the age (post-conceptional days) of all samples in the human PFC dataset and then trained a regularized linear model by regressing the transformed ages against the riBAQ values of all PSD proteins. Specifically, ridge regression was performed using the R package glmnet ^89^ with λ (equals 0.4466836) selected by ten-fold cross-validation. The equivalent human PSD ages for macaque and mouse samples were then predicted using the trained model and listed in Supplementary Table 11b.

The predicted PSD ages of human, macaque, and mouse samples were regressed against the real post-conceptional ages using a linear regression model to obtain the slope coefficients as an estimator of the PSD maturation rate. For the macaque samples, because the PSD maturation rate appears to be different before and after the age of one year, the predicted PSD ages were regressed against the real post-conceptional ages using a linear spline model with the knot set at post-conceptional day 330 using the R package lspline.

Similar predictions were made based on the human V1 dataset. For ridge regression, λ (equals 0.7943282) was selected by ten-fold cross-validation. The equivalent human PSD ages based on the human V1 dataset were listed in Supplementary Table 11c.

### Primary neuronal culture

Primary human cortical neurons were prepared from GW21 to GW23 human dorsal cortical tissue samples. The cortical plate and subplate were dissected and dissociated using the Papain Dissociation System (Worthington Biochemical). Dissociated neurons were resuspended in a plating medium (Neurobasal medium supplemented with 1xB27, 2 mM GlutaMAX, and antibiotics) and plated into tissue culture plates coated with PEI-laminin or containing a 12 mm coverslip pre-coated with PDL and laminin (Corning 354087) at the density of 100K cells/cm^2^. Cells were cultured in a humidified incubator with 5% CO_2_ and 8% O_2_. On days in vitro (DIV) 1, the medium was changed to maturation medium (BrainPhys medium supplemented with 1 × B27, 1 × N2, 20 ng/mL BDNF, 20 ng/mL GDNF, 1mM dibutyryl-cAMP, 200 nM ascorbic acid, 1 μg/mL laminin, and antibiotics). Half of the medium was changed with fresh medium every 3–4 days until harvest.

Primary mouse cortical neurons were prepared from postnatal day (P) 0 C57BL6/J mice. The neocortices were dissected, dissociated, and plated using the same procedure as primary human neurons.

### Synaptosome preparation from cultured primary neurons

About two million neurons were harvested at the indicated DIV in 2 mL homogenization buffer (0.32 M sucrose, 2 mM EDTA, and 4 mM HEPES, pH 7.4) with freshly added protease and phosphatase inhibitors (Roche) and homogenized using a 10 mL tissue grinder. Postnuclear supernatants were obtained via 1,000 × g spin. The supernatant (S1) was spun at 10,000 × g for 15 min, and the pelleted membranes were resuspended in 1 mL 0.8M sucrose buffer and spun at 20,000 × g for 30 min. The pellet (synaptosomes) was resuspended in the homogenization buffer and stored at −80 °C before Western blot analysis. Protein concentrations were determined using the BCA assay (Pierce).

### Plasmid cloning

pmCherry-1 (632525, Clontech) was used as the backbone and control vector for the overexpression experiments. mCherry-ARHGEF7 was a gift from Dorus Gadella (Addgene # 129611). The cDNA of RASGRF2 was cloned into pmCherry-1 from R777-E241 Hs.RASGRF2, a gift from Dominic Esposito (Addgene # 70525). pLKO-RFP-shCntrl, a gift from William Kaelin (Addgene # 69040), was used as the backbone and control vector for the knockdown experiments. Sequences of shRNAs against *ARHGAP23* or *SRGAP1* can be found in Supplementary Table 12a.

### Validation of shRNA knockdown efficiency

HEK293T (ATCC CRL-3216) cultured in Dulbecco’s Modified Eagle Medium (DMEM) (Corning) containing 10% Fetal Bovine Serum (Hyclone) and antibiotics (Penicillin/Streptomycin) were plated in 12-well plates. The next day, 1 μg of corresponding shRNA vectors were transfected into the cells using Lipofectamine 3000 (Invitrogen). Cells were harvested 48 hours after transfection for mRNA extraction using the RNAeasy mini plus kit (Qiagen). RNA quantity and quality were checked using NanoDrop 1000 (Thermo Scientific). qRT-PCR was performed using the ViiA 7 Real-time PCR System with PowerUp SYBR Green Master Mix and analyzed with comparative Ct method normalized against the housekeeping gene *GAPDH*. Primers used were listed in Supplementary Table 12b.

### Plasmid DNA transfection into primary neurons and immunocytochemistry

Human neurons were transfected on DIV28 with pEGFP-C1 (0.3 μg/well, Clontech) plus vectors expressing mCherry, mCherry-ARHGEF7, mCherry-RASGRF2, tRFP-shControl, tRFP-sh*ARHGAP23*-1, tRFP-sh*ARHGAP23*-2, tRFP-sh*SRGAP1*-1, or tRFP-sh*SRGAP1*-2 (0.7 μg/well) with lipofectamine 2000. On DIV42, human neurons were fixed with 4% formaldehyde/4% sucrose in PBS and permeabilized/blocked with PBS-based blocking buffer containing 10% donkey serum, 0.2% gelatin, and 0.1% Triton X-100 at room temperature for 1 h. Samples were then incubated with primary antibodies at 4 °C overnight. The next day, samples were washed in PBS three times and incubated with secondary antibodies in the blocking buffer at room temperature for 1 h. Samples were then washed in PBS twice, counterstained with DAPI, and washed in PBS once more. Z-stack images were acquired with a Leica TCS SP8 using a 63× oil immersion objective. Dendritic spine analysis was performed using Imaris (Oxford Instruments). Spine density and morphology were measured from secondary or tertiary dendrites. Automatic spine classification was done using the “Classify Spines Xtension” in Imaris with the following rules: Stubby—length(spine)<1 μm; Mushroom—length(spine)<3 μm and max_width(head) > mean_width(neck)*2; Long Thin—length(spine)<3 μm but not Mushroom; Filopodia— length(spine)>=3 μm. Mouse neurons were transfected on DIV5 and analyzed on DIV8 in a similar manner.

For surface GRIA1 staining, after fixation, neurons were blocked by PBS-based blocking buffer containing 10% donkey serum and 0.2% gelatin at room temperature for 1 h. Samples were then incubated with primary antibodies against extracellular GRIA1 under the non-permeabilized condition at room temperature for 1 h. Next, samples were washed in PBS three times and permeabilized/blocked with blocking buffer containing 0.1% Triton X-100 at room temperature for 1 h. Samples were then incubated with primary antibodies against intracellular proteins at 4 °C overnight. The rest steps of the procedure were the same as mentioned above.

The following antibodies were used: GFP (Aves, GFP-1020, 1:1000), mCherry (Invitrogen, M11217, 1:500), SYN1 (Cell Signaling Technology, 5297S, 1:250), DLG4 (Synaptic Systems, N3702-AF647-L, 1:250), tRFP (OriGene, TA150061, 1:250), GRIA1 (Alomone, AGC-004, 1:25).

### Whole-cell recording of primary human cortical neurons

Human neurons transfected with pEGFP-C1 plus vectors expressing mCherry, mCherry-ARHGEF7, or mCherry-RASGRF2 were recorded between DIV42 and DIV49. Whole-cell recordings were performed in the maturation medium supplemented with 1 mM TTX at room temperature. The voltage was set at −60 mV. Recording pipettes (4-10 mOhm) were filled with the intrapipette solution containing 140 mM K-gluconate, 2 mM MgCl_2_, 10 mM HEPES, 0.2 mM EGTA, 4 mM MgATP, 0.3 mM NaGTP, 10 mM Phosphocreatine d-tris, and 0.25% biocytin. Signals were collected at a sampling rate of 10K, using a 10K Bessel filter (MultiClamp 700B, Axon Instruments, Molecular Devices) and digitized (Digidata, Axon Instruments, Molecular Devices). Putative mEPSC events were extracted in the open-source software Stimfit 0.15.8 (https://github.com/neurodroid/stimfit).

### MAGMA analysis of PSD modules using GWAS data of human cognitive functions

MAGMA v1.09 ^90^ was used to determine whether human PSD modules are enriched for common variants associated with human cognitive function.

GWAS summary statistics for human cognitive function studies conducted by the UK Biobank were downloaded from the GWAS ATLAS resource (https://atlas.ctglab.nl/) ^91^. We analyzed GWAS data of Reaction Time Test for processing speed, Fluid Intelligence Test for Fluid intelligence, Trail Making Test Part B for executive function, Pairs Matching Test for visual declarative memory, Numeric Memory Test for working memory, and Prospective Memory Test for prospective memory.

For gene analysis in MAGMA, the 1,000 Genomes European panel was used to estimate LD between SNPs, and the SNP-wise Mean model was used as the gene analysis model. For gene-set analysis, several common technical confounders are included in the linear regression model as covariates. These confounders include gene size, gene density, the inverse of the mean minor allele count of variants, and their log-transformed values. The resulting nominal p-values were adjusted using the Benjamini-Hochberg method ^74^. The detailed results can be found in Supplementary Table 13a.

### Module enrichment of activity-dependent proteins

Lists of proteins whose abundances depend on neuronal activities were obtained from Schanzenbächer et al., 2018 ^48^. The rat protein list was converted to human genes using ortholog data obtained from the Alliance of Genome Resources (https://www.alliancegenome.org/). The one-sided hypergeometric test was used to determine if there was a significant overlap with individual PSD modules. The union of genes encoding PSD proteins and those expressed in the human neocortex was used as the background. The resulting nominal p-values were adjusted using the Benjamini-Hochberg method. The detailed results can be found in Supplementary Table 13b.

### Gene constraint analysis

LOEUF scores, missense Z-scores, and synonymous Z-scores for all human genes were downloaded from gnomAD v2.1.1 (gnomad.broadinstitute.org/) ^92^. The union of genes encoding PSD proteins and those expressed in the human neocortex was used as the background. Data were plotted using the geom_boxplot() function from the R package ggplot2 with default settings for elements. The Kruskal-Wallis rank sum test with *post hoc* Dunn’s test was used to compare differences in scores between each category of genes.

### Module enrichment of de novo variants associated with neurodevelopmental disorders

*De novo* variants in neurodevelopmental disorders, including epilepsy, developmental delay, autism spectrum disorder, intellectual disability, and schizophrenia, were obtained from the denovo-db v.1.6.1 (https://denovo-db.gs.washington.edu/denovo-db/) ^93^. We also included one additional epilepsy dataset ^94^ and two additional schizophrenia datasets ^95,96^ not included in the denovo-db database. We defined “stop-gained”, “start-lost”, “stop-gained-near-splice”, “frameshift”, “frameshift-near-splice”, “splice-donor”, and “splice-acceptor” mutations as protein-truncating variants (PTVs) and “missense” and “missense-near-splice” mutations as missense variants. The number of PTVs or missense variants for a gene in a disorder was defined as the number of individuals with the disorder harboring PTVs or missense variants in the gene. The one-sided Fisher’s exact test was used to determine if there is a significant enrichment of *de novo* variants in genes within individual PSD modules compared with those outside the module. The resulting nominal p-values were adjusted using the Benjamini-Hochberg method. The summarized results can be found in Supplementary Table 13c.

### MAGMA analysis of PSD modules using GWAS data of human psychiatric disorders

GWAS summary statistics for schizophrenia (June 2018 release), bipolar disorder (June 2018 release), autism spectrum disorder (November 2017 release), major depressive disorder (mdd2019edinburgh), and attention-deficit/hyperactivity disorder (January 2022 release) were downloaded from The Psychiatric Genomics Consortium database (https://www.med.unc.edu/pgc/) ^97–100^. MAGMA analysis was done in the same way as described above. The detailed results can be found in Supplementary Table 13d.

### Module enrichment of misexpressed genes after the onset of psychiatric disorders

Gene expression data from brain samples in schizophrenia, bipolar disorder, autism spectrum disorder, major depressive disorder, and controls were obtained from Gandal et al., 2018 ^101^. Genes with an adjusted p-value less than 0.05 were considered misexpressed genes in a psychiatric disorder. The summary can be found in Supplementary Table 13e. The one-sided hypergeometric test was used to determine if there was a significant overlap with individual PSD modules. The union of genes encoding PSD proteins and those expressed in the human neocortex was used as the background. The resulting nominal p-values were adjusted using the Benjamini-Hochberg method.

## Code availability

Code used for data analysis in this manuscript can be found at https://github.com/alexwang1001/PSD_development/.

## Data and materials availability

All raw proteomic data were deposited to ProteomeXchange through MassIVE (human PFC dataset: MSV000091887, human V1 dataset: MSV000091888, Macaque dataset: MSV000091889, Mouse dataset: MSV000091890). All processed data are available in the auxiliary supplementary tables and at an online interactive portal (https://liwang.shinyapps.io/PSD_development_explorer/).

## Acknowledgments

We thank NIH NeuroBioBank, the University of Maryland School of Medicine Brain and Tissue Bank for providing post-mortem brain tissue samples. This work used the Vincent J. Proteomics/Mass Spectrometry Laboratory at UC Berkeley, supported in part by NIH S10 Instrumentation Grant S10RR025622. We thank Lori Kohlstaedt for her help in mass spectrometry data analysis. We thank Stephan Sanders, Roger Nicoll, Neelroop Parikshak, Tanzila Mukhtar, Mengyi Song, and Huda Zoghbi for comments on the manuscript. This study was supported by National Institute of Neurological Disorders and Stroke grant 5R35NS097305 to A.R.K., National Human Genome Research Institute grant HG010898 to N.S., and National Institute of Mental Health grant MH124619 to N.S..

## Author contributions

Conceptualization: L.W.; data curation: L.W.; formal analysis: L.W.; funding acquisition: A.R.K.; investigation: L.W., L.Z., A.C.-S., S.G.-G., S.W., M.W., B.H., T.L.; methodology: L.W., K.P., L.Z., A.C.-S., S.G.-G., Jiani Li, P.L., Jingjing Li, J.M.A.-G.; resources: S.W., Y.P., E.J.H., E.A.W., M.F.P., R.K., N.S., A.P.; software: L.W., K.P.; supervision: X.P., J.M.A.-G., A.A.-B., Z.L., A.R.K.; visualization: L.W., K.P.; writing – original draft: L.W.; writing – review & editing: all authors.

## Supplementary Materials

Supplementary Figure 1

Supplementary Tables 1 to 13

**Extended Data Fig. 1.**
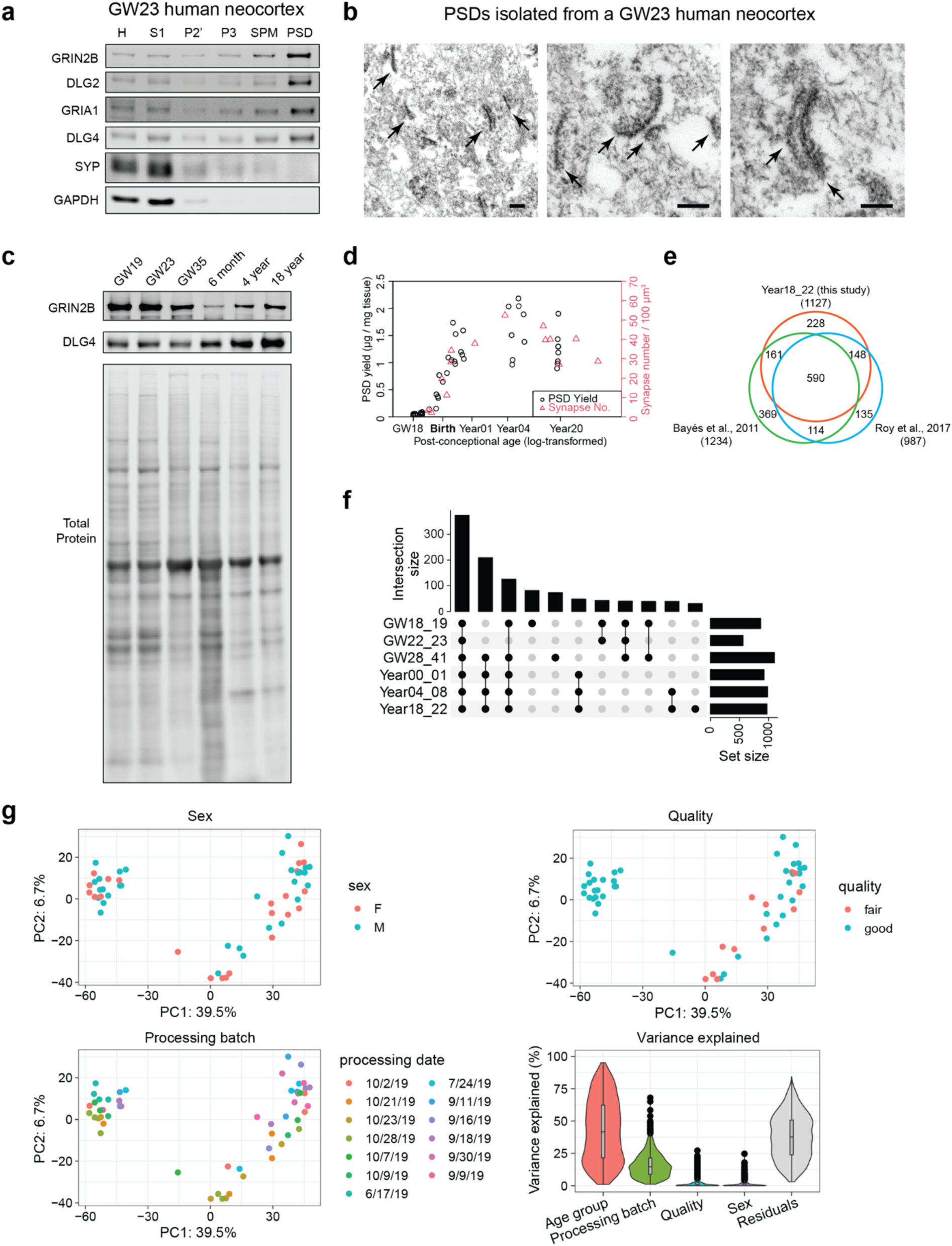
Isolation of PSD from immature and mature human cortices. **a**, Western blot analysis of different subcellular fractions of a GW23 sample demonstrating enrichment of PSD proteins and depletion of presynaptic SYP and cytoplasmic GAPDH in the PSD fraction. **b**, Electron micrographs of the PSD fraction isolated from a GW23 sample (scale bar: 200 nm). Arrows denote structures resembling the PSD. **c**, Western blot analysis of purified PSDs from different age groups demonstrating changes in GRIN2B and DLG4 during development. **d**, Correlation between PSD yield and synapse number of the human prefrontal cortex. **e**, Venn diagram showing the overlap between Year18_22 samples in this study and the human PSD proteomes published in Roy et al., 2017 and Bayés et al., 2011. **f**, UpSet plot describing the number of identified proteins and their overlaps at each age group. **g**, PCA plots of the samples colored by various covariates and variance explained by individual covariates.

**Extended Data Fig. 2.**
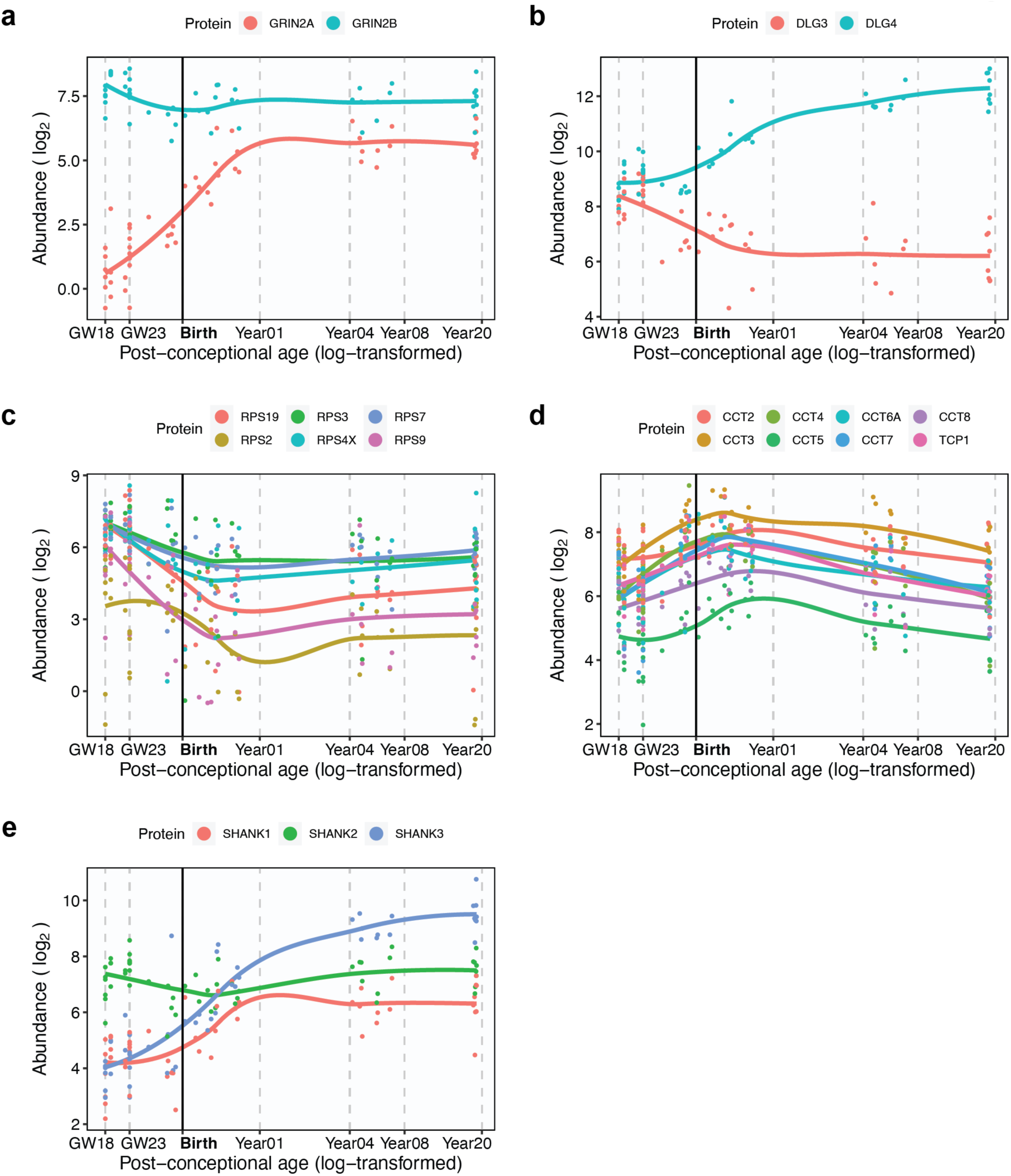
Examples of PSD protein abundance patterns in human cortical development. **a**, Abundance patterns of GRIN2A and GRIN2B. **b**, Abundance patterns of DLG3 and DLG4. **c**, Abundance patterns of the 40S ribosomal subunits. **d**, Abundance patterns of the TRiC subunits. **e**, Abundance patterns of SHANK family scaffolding proteins.

**Extended Data Fig. 3.**
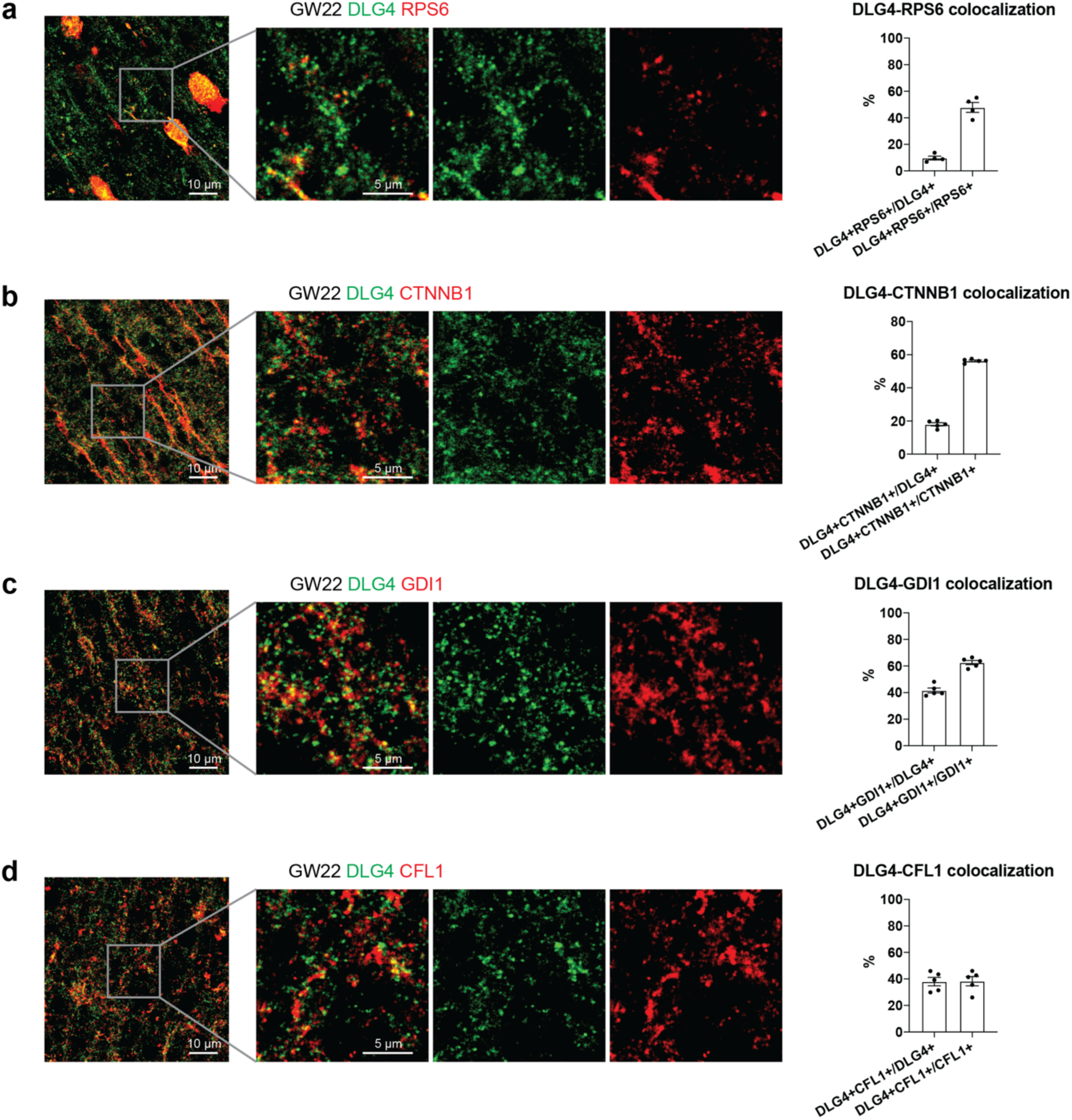
Colocalization of early PSD proteins with DLG4 in immature human neocortex. **a–d**, Colocalization of RPS6 (**a**), CTNNB1 (**b**), GDI1 (**c**), or CFL1 (**d**) with DLG4 in GW22 human neocortex (scale bar: 10 μm or 5 μm as indicated in the figure).

**Extended Data Fig. 4.**
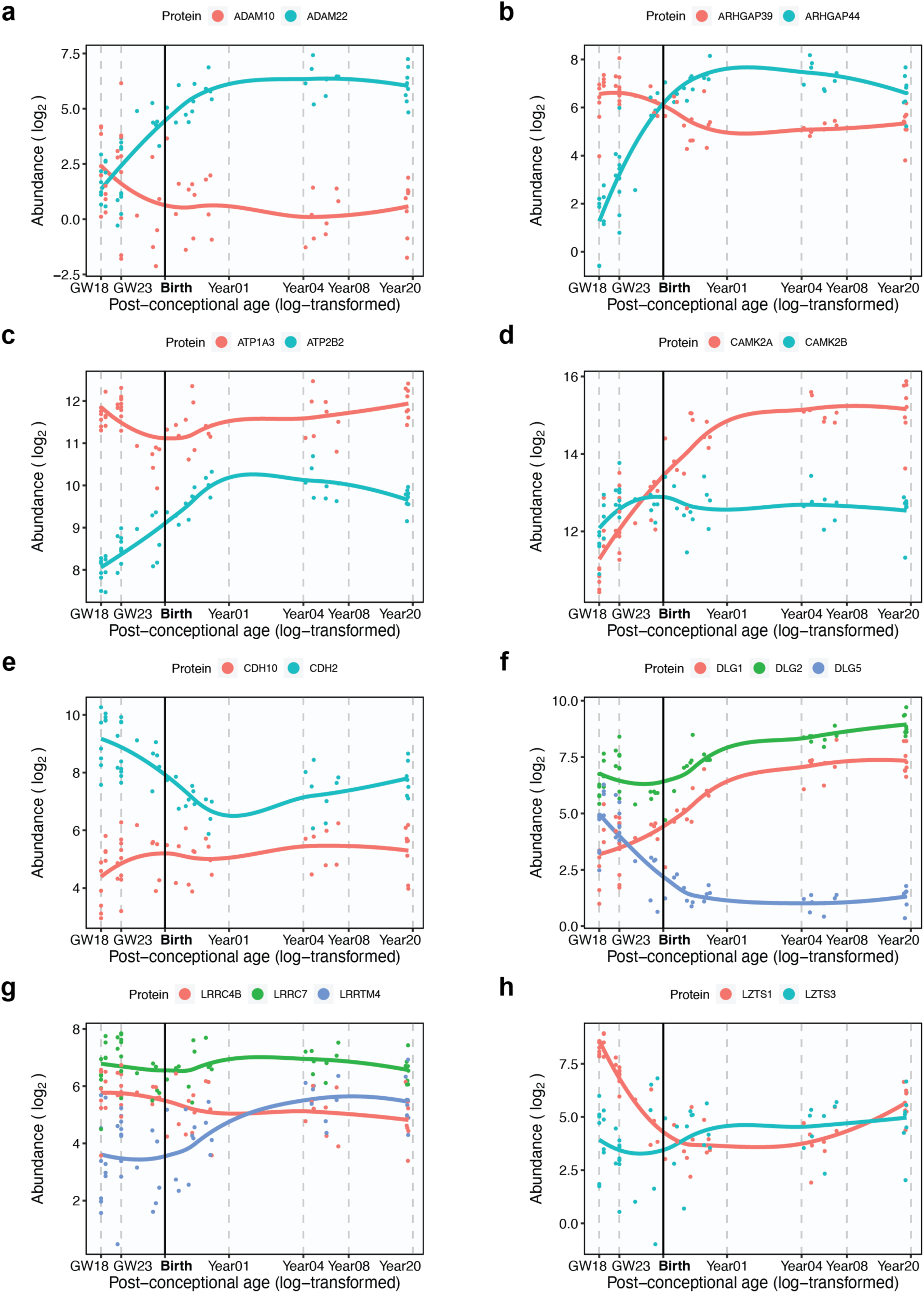
PSD protein paralogs subjected to reciprocal developmental changes. **a–h** Examples of PSD protein paralogs identified in this study that undergo reciprocal developmental changes.

**Extended Data Fig. 5.**
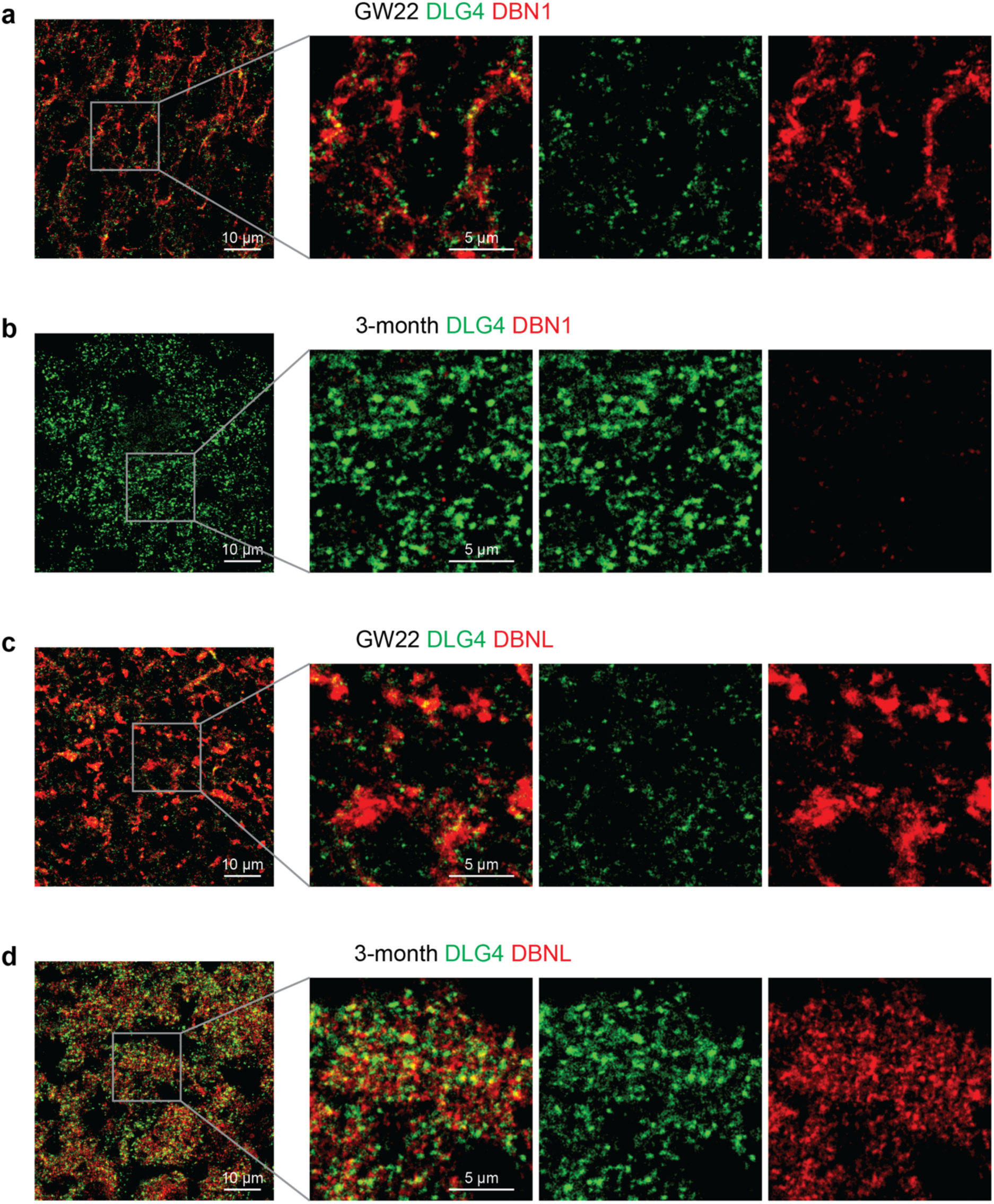
Quantification of DBN1 and DBNL in the PSD of the human neocortex. **a–d**, Colocalization of DBN1 (**a** and **b**) and DBNL (**c** and **d**) with DLG4 in the GW22 (**a** and **c**) and 4-month (**b** and **d**) human neocortex (scale bar: 10 μm or 5 μm as indicated in the figure).

**Extended Data Fig. 6.**
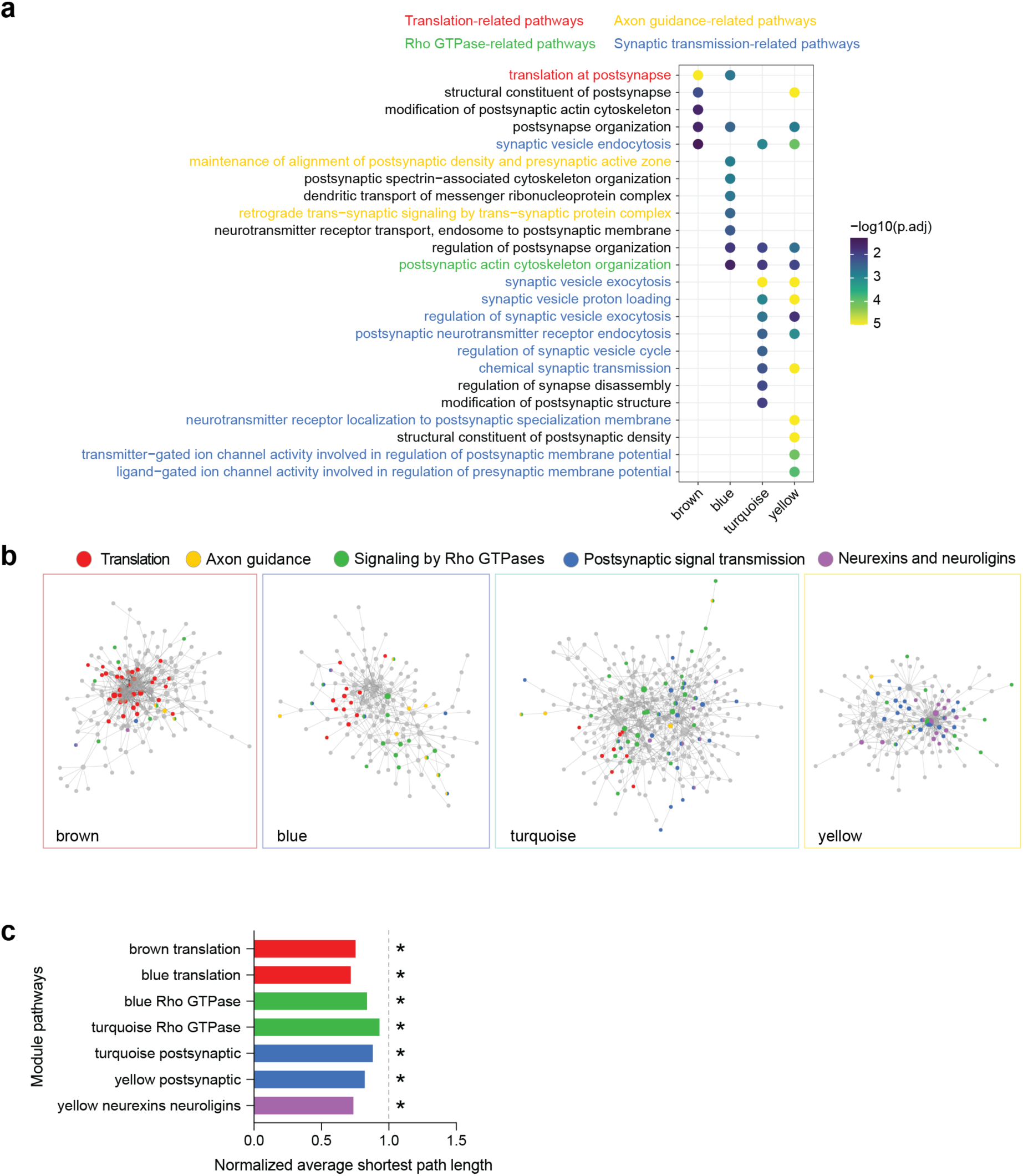
Pathway enrichment in individual PSD modules. **a**, SynGO biological pathway enrichment analysis of each module. **b**, PPI-co-abundance network of each module highlighting proteins in enriched pathways. **c**, The normalized average shortest path lengths of pathways in individual modules. The asterisks denote that the average shortest path length is significantly shorter within pathway proteins than between pathway and non-pathway proteins. *p < 0.01; Wilcoxon rank sum test.

**Extended Data Fig. 7.**
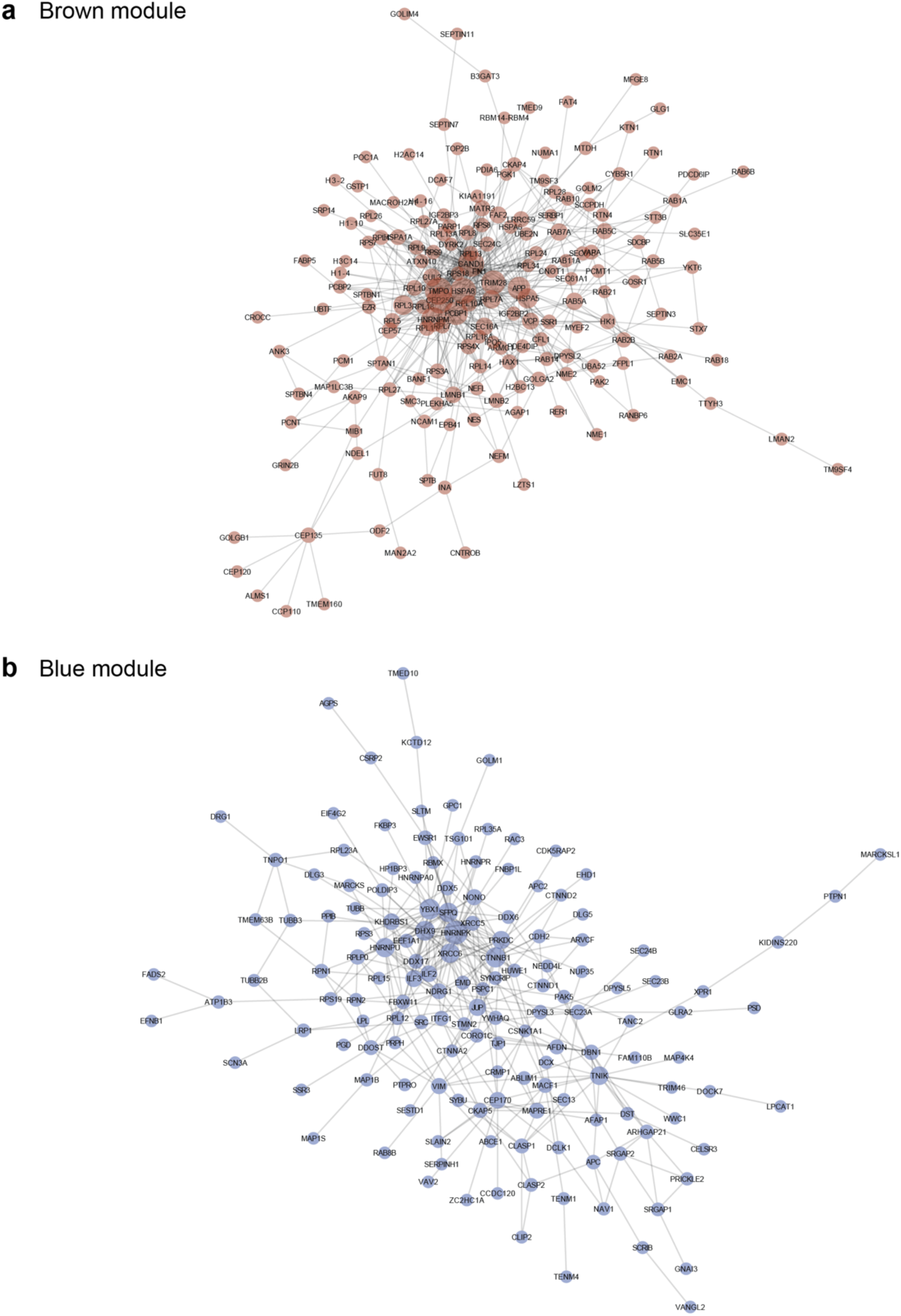
PPI-co-abundance network of the brown (a) and the blue (b) modules.

**Extended Data Fig. 8.**
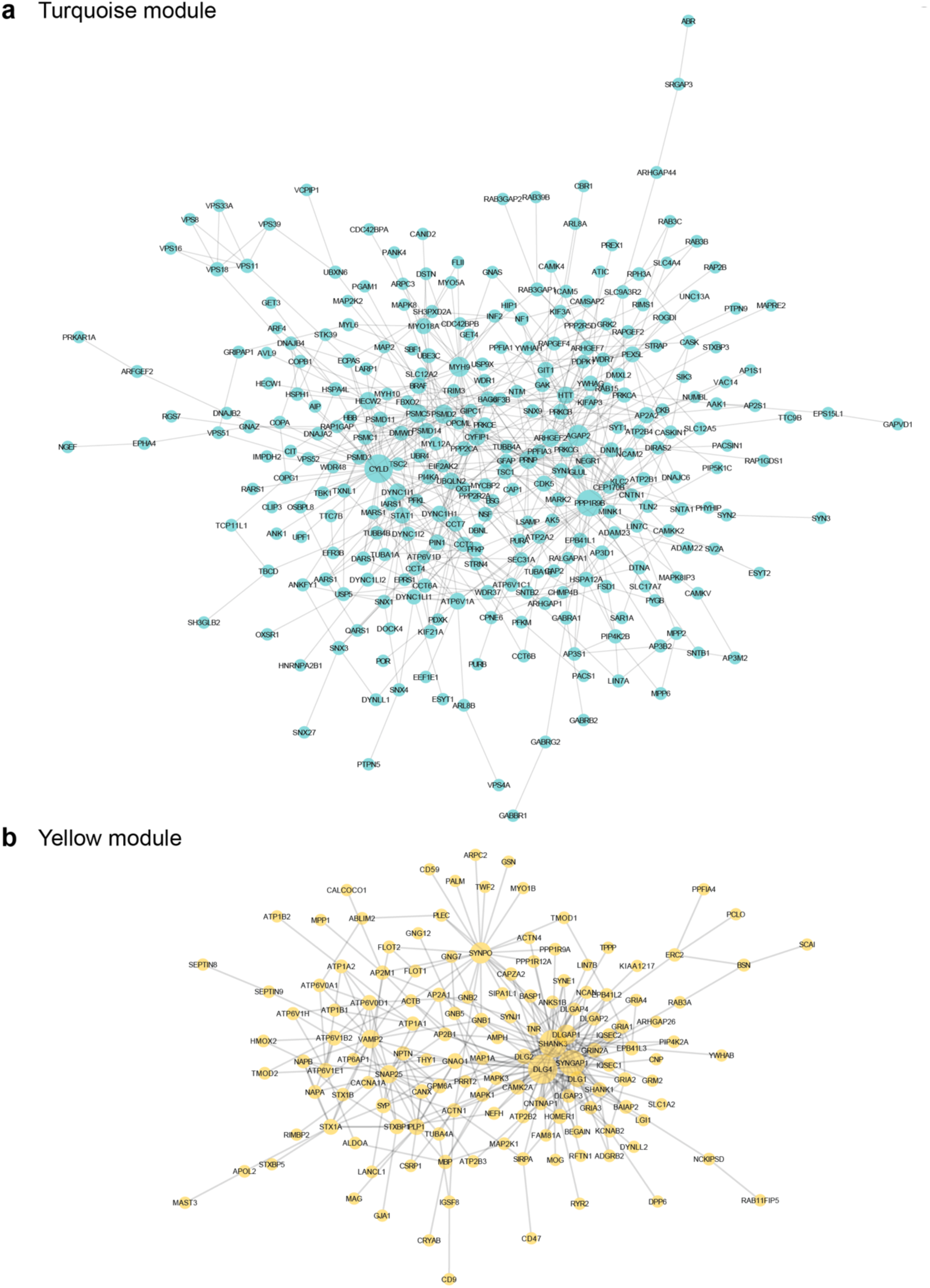
PPI-co-abundance network of the turquoise (a) and the yellow (b) modules.

**Extended Data Fig. 9.**
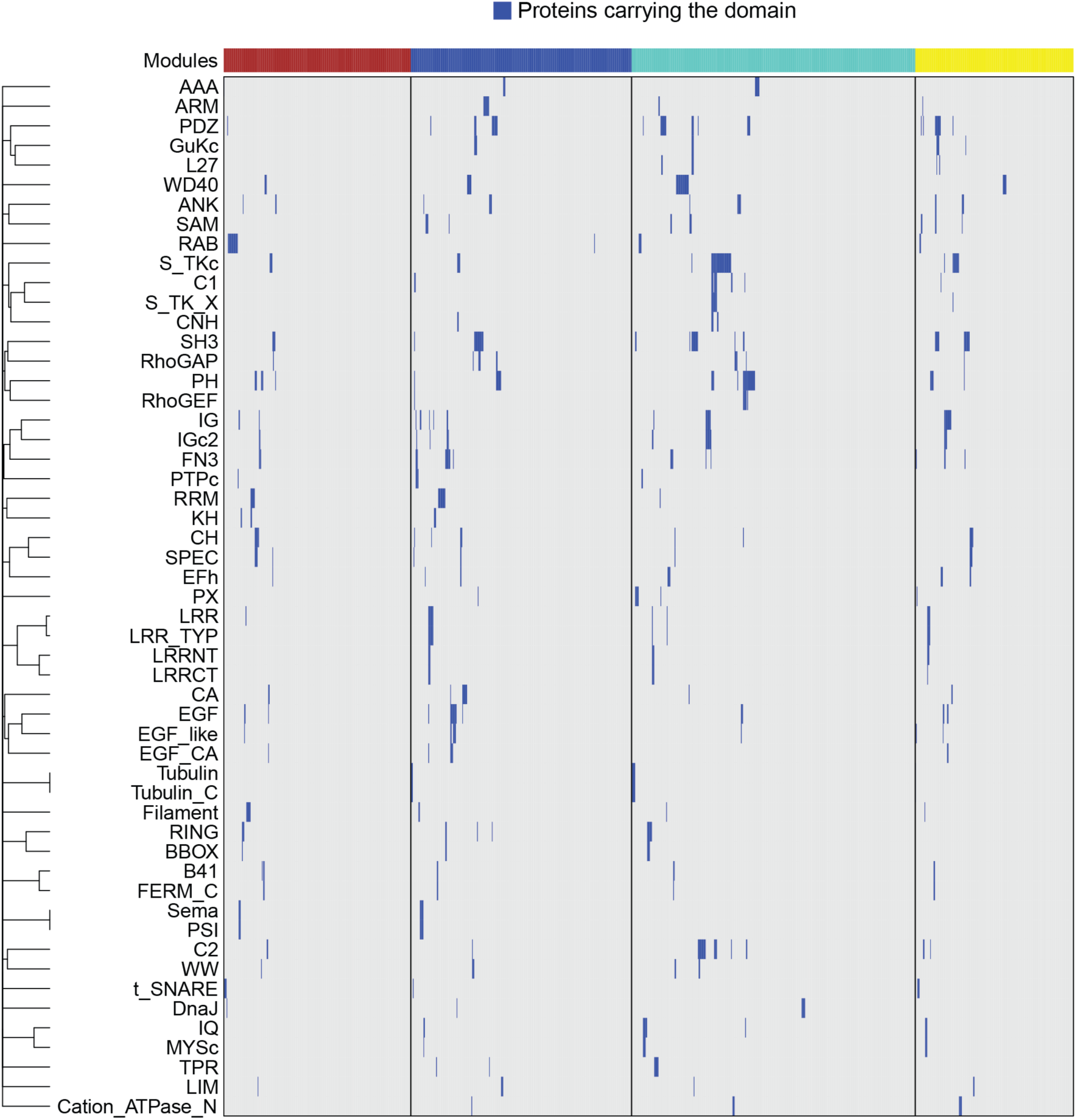
Protein domains in individual PSD proteins. The rows are clustered based on the Jaccard distance.

**Extended Data Fig. 10.**
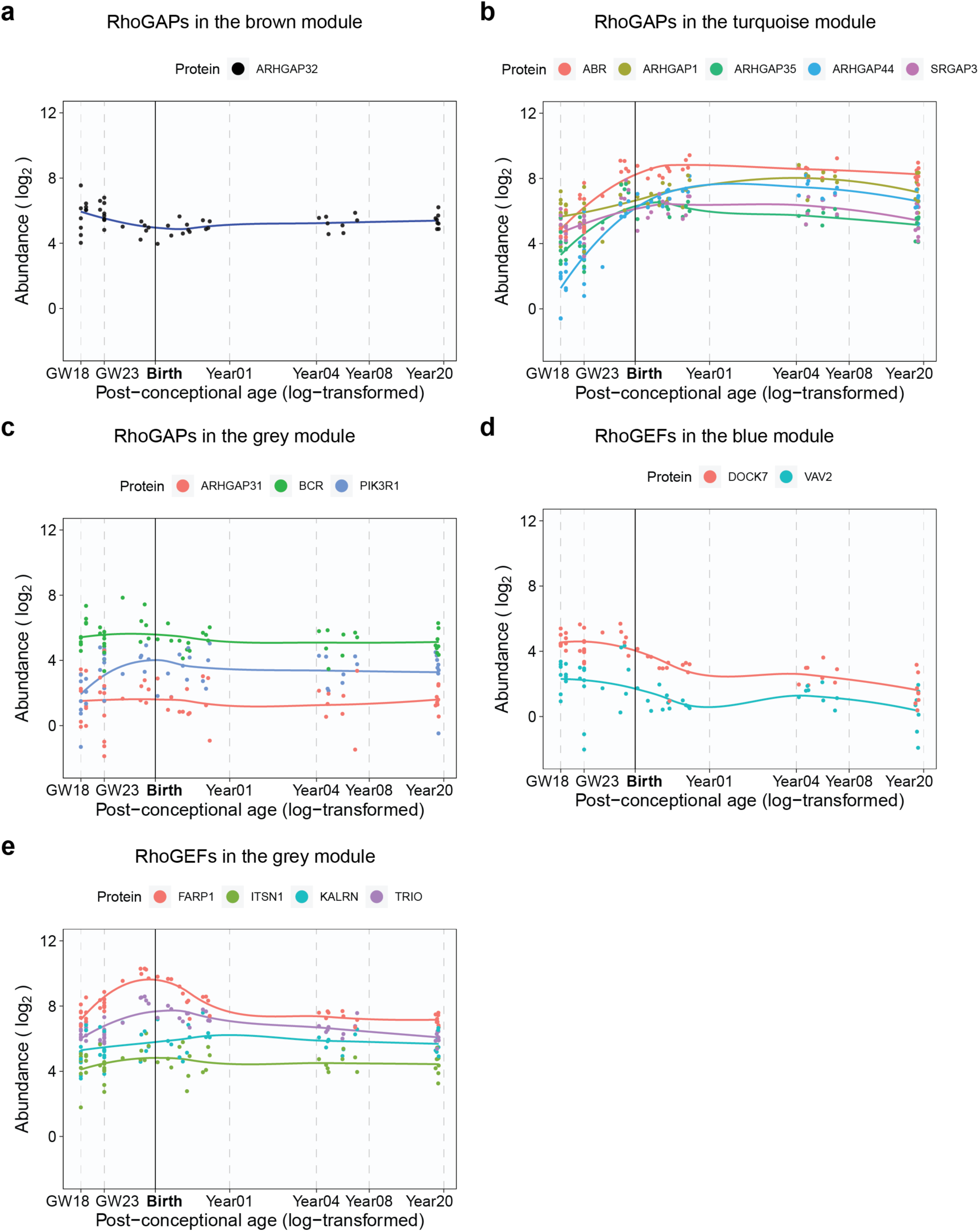
Abundance patterns of RhoGAPs and RhoGEFs not listed in Fig. 2f.

**Extended Data Fig. 11.**
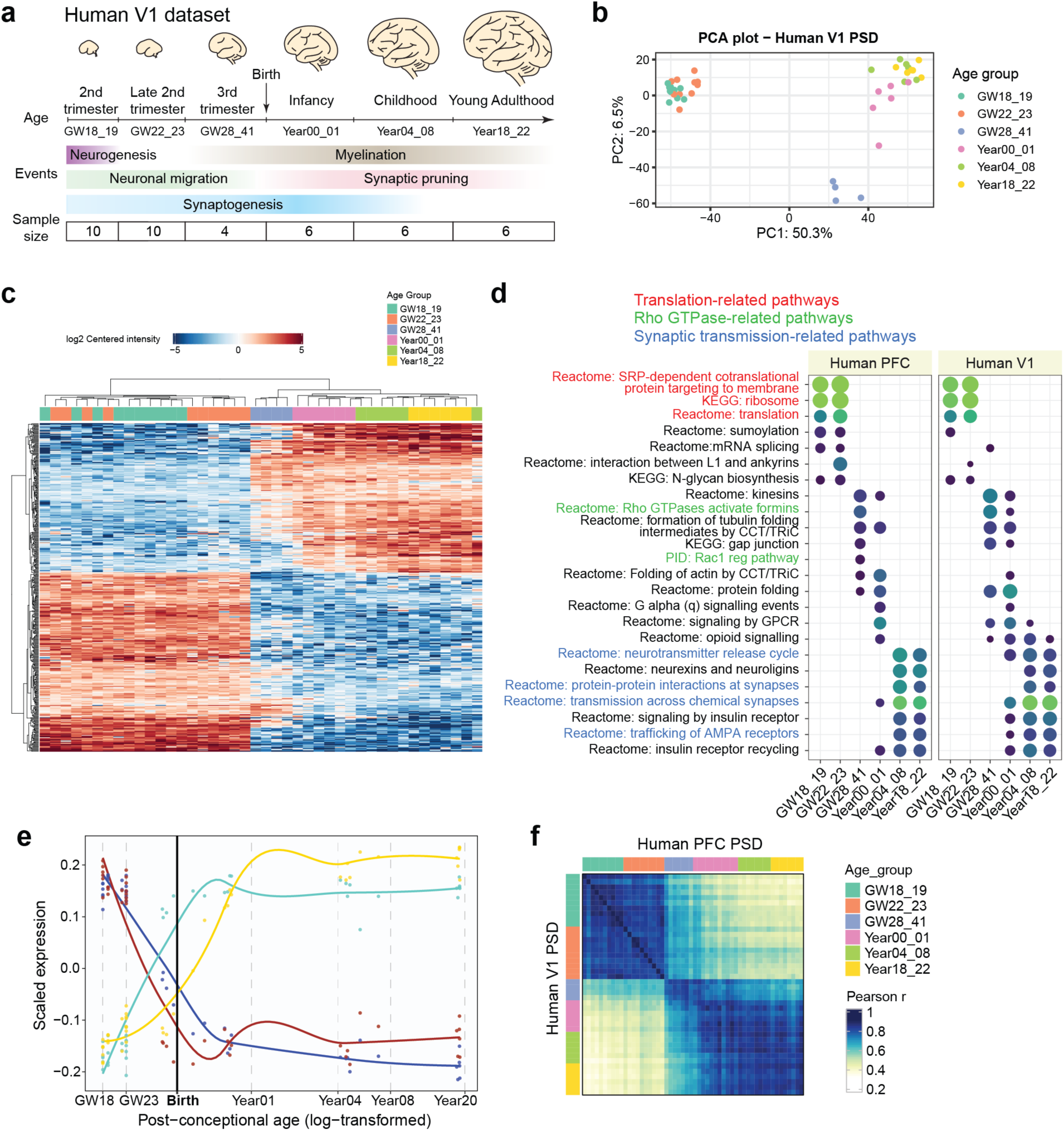
Changes in PSD composition during human V1 development. **a**, Schematic illustrating the developmental stages of samples in the human V1 dataset. **b**, PCA plots of samples in the human V1 dataset colored by their age groups. **c**, Hierarchical clustering of the samples in the human V1 dataset based on proteins with differential abundance. **d**, Gene set enrichment analysis for individual age groups in the human PFC and V1 dataset. NES: normalized enrichment score. **e**, Scaled abundance patterns (module eigengene values) of four protein modules in the human V1 dataset. **f**, Similarity matrices representing pairwise Pearson correlations between human PFC and human V1 samples.

**Extended Data Fig. 12.**
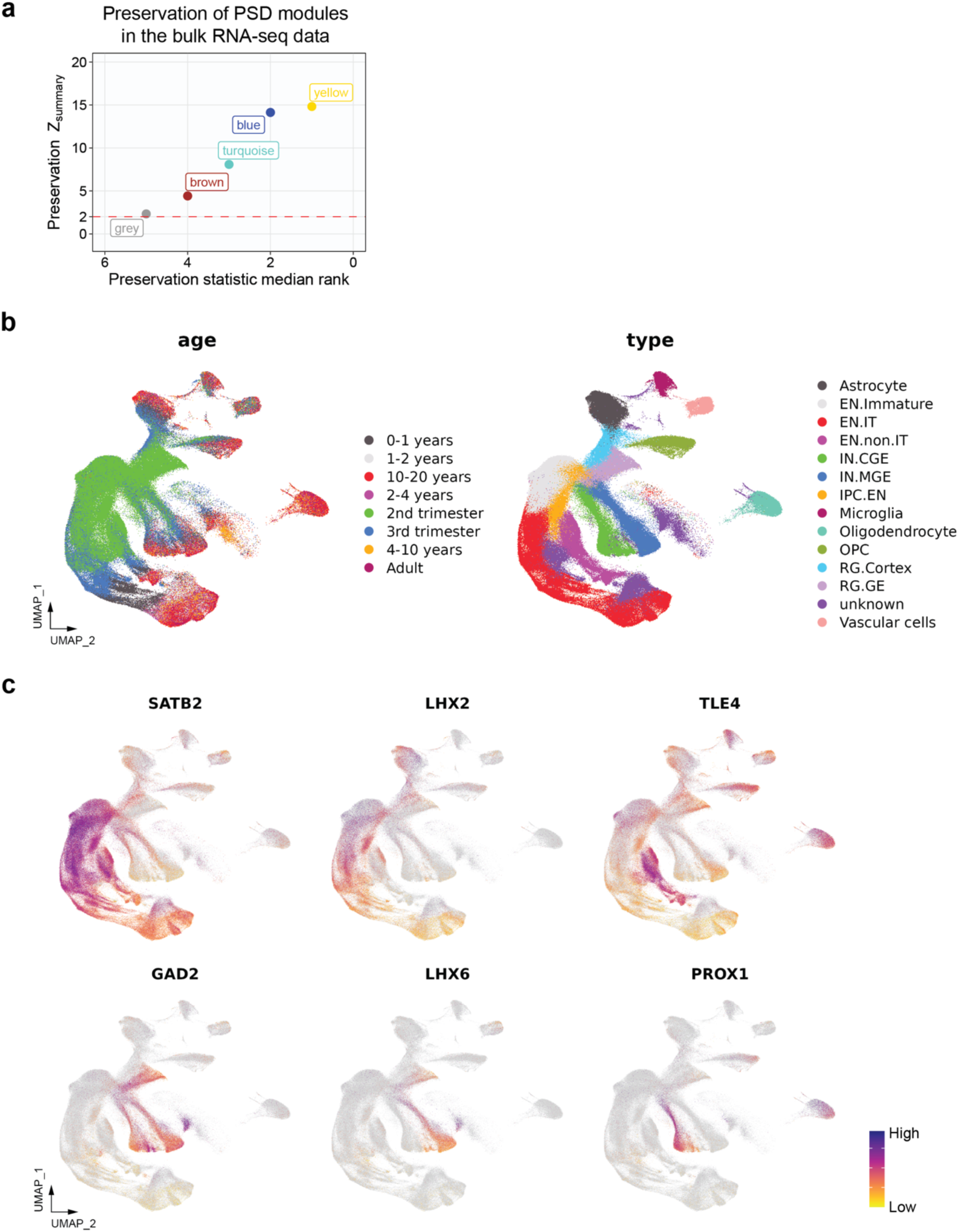
Preservation of human PSD modules at the RNA level and overview of the single-nucleus RNA-seq data from the developing human neocortex. **a**, Preservation of Human PSD modules in the bulk RNA-seq data. **b**, UMAP plots showing the distribution of age groups and cell types in the single-nucleus RNA-seq data from the developing human neocortex. **c**, UMAP plots showing the expression patterns of neuronal subtype-specific markers in the single-nucleus RNA-seq data from the developing human neocortex.

**Extended Data Fig. 13.**
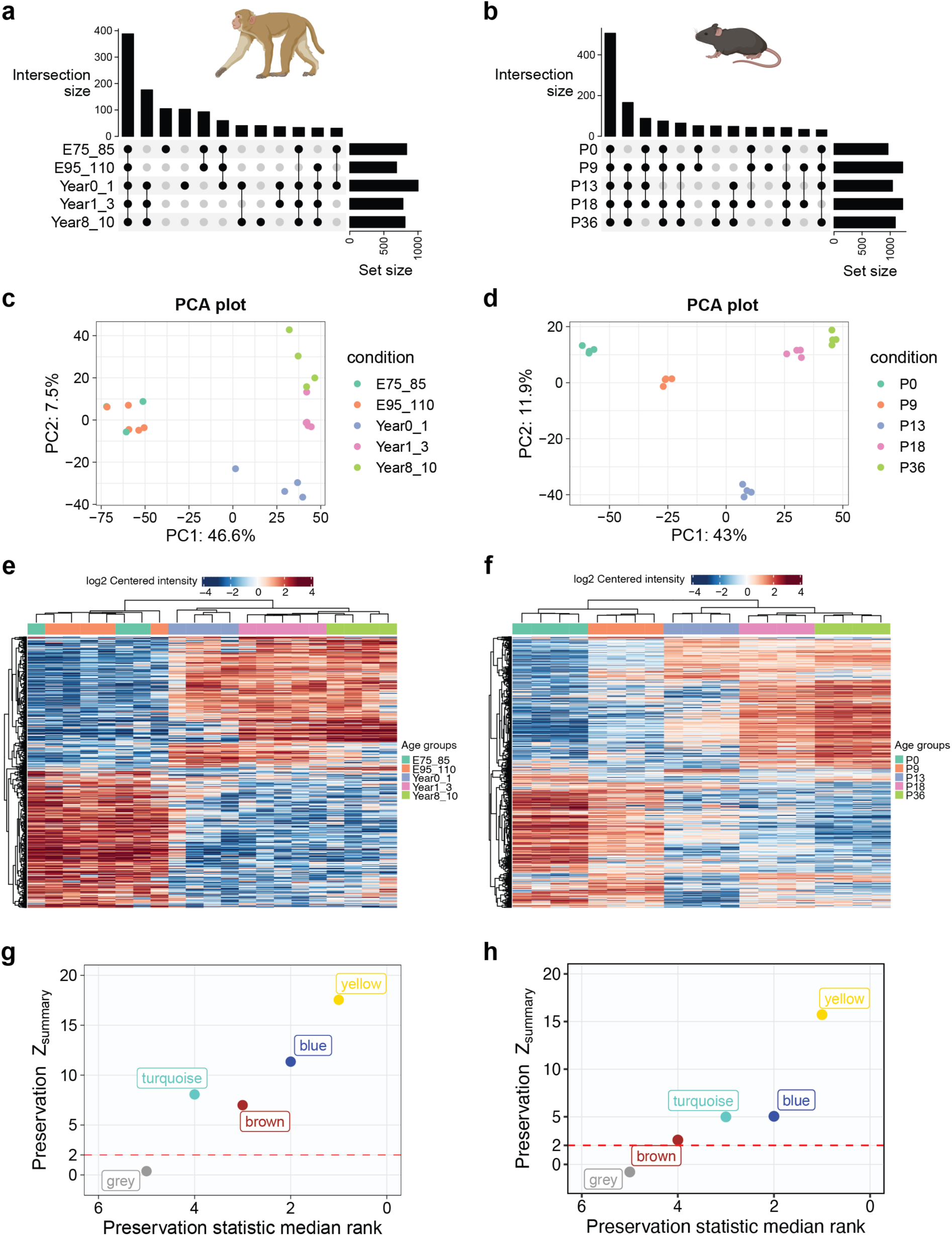
Changes in PSD composition during macaque and mouse neocortical development. **a**, UpSet plot describing the number of identified proteins and their overlaps at each age group of the macaque dataset. **b**, UpSet plot describing the number of identified proteins and their overlaps at each age group of the mouse dataset. **c**, PCA plots of the macaque samples colored by various covariates. **d**, PCA plots of the mouse samples colored by various covariates. **e**, Hierarchical clustering of the macaque samples based on proteins with differential abundance. **f**, Hierarchical clustering of the mouse samples based on proteins with differential abundance. **g**, Preservation of Human PSD modules in the macaque PSD proteomic data. **h**, Preservation of Human PSD modules in the mouse PSD proteomic data.

**Extended Data Fig. 14.**
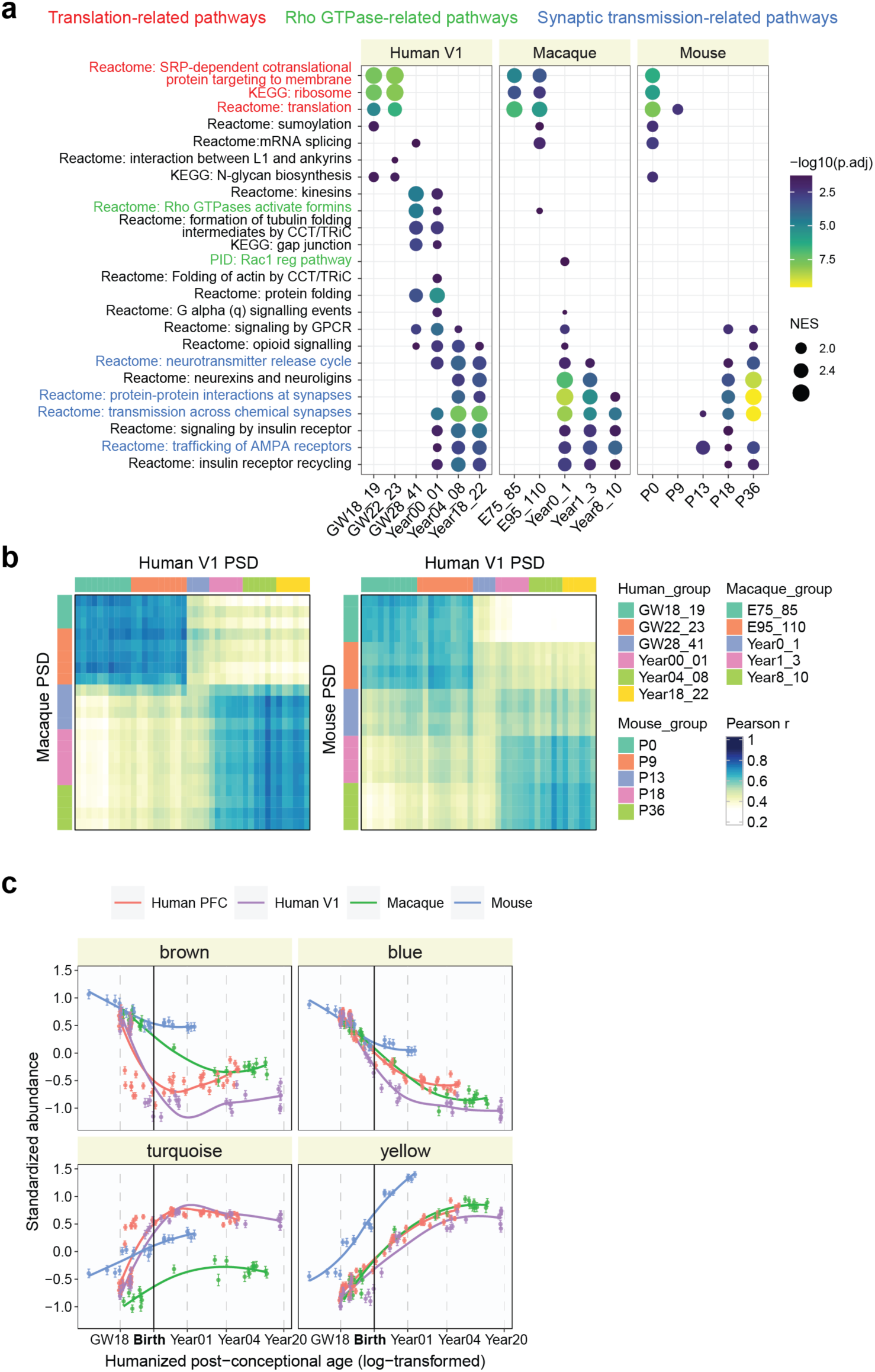
Comparison of PSD development across the human V1, macaque, and mouse datasets. **a**, Gene set enrichment analysis for individual age groups across datasets. NES: normalized enrichment score. **b**, Similarity matrices representing pairwise Pearson correlations between human V1, macaque, and mouse samples. **c**, Standardized abundance patterns of proteins in the four PSD modules across regions and species along the humanized age based on the human V1 dataset.

**Extended Data Fig. 15.**
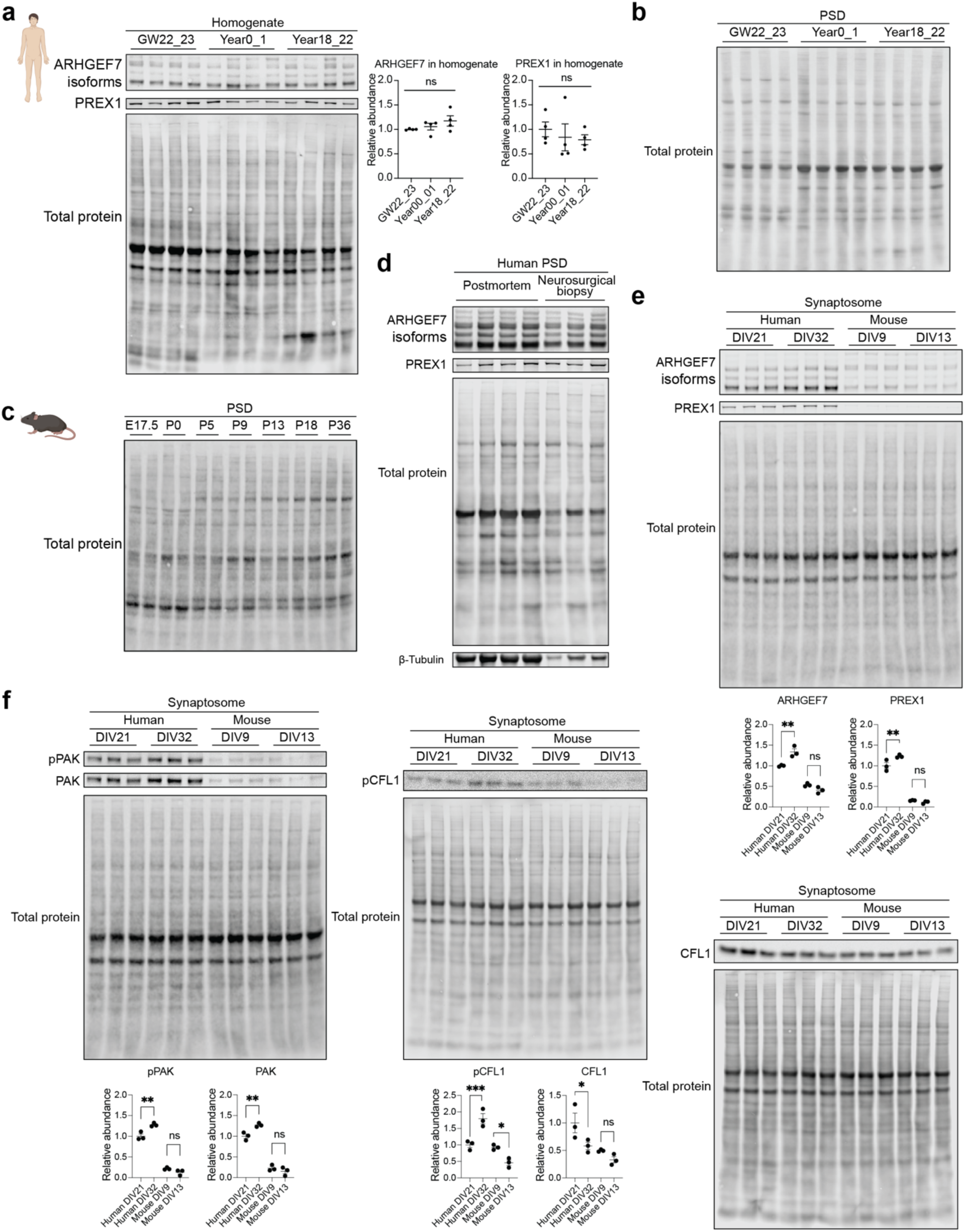
RhoGEF levels and activities during synapse development. **a**, Immunoblots and quantification of representative RhoGEFs in the whole homogenate of the developing human (n = 4, 4, 4 samples) and mouse (n = 2, 2, 2, 2, 2, 2, 2 samples) PSD. One-way ANOVA with Holm-Sidak’s multiple comparisons test. **b**, Total protein staining for blots of the human PSD in Fig. 5b. **c**, Total protein staining for blots of the mouse PSD in Fig. 5b. **d**, Comparison of adult PSD samples from postmortem brain tissues and neurosurgical biopsy tissues by western blot analysis. **e**, Immunoblots and quantification of representative RhoGEFs in the synaptosomes of cultured primary human (n = 3, 3 samples) and mouse cortical neurons (n = 3, 3 samples). **p < 0.01; one-way ANOVA with Holm-Sidak’s multiple comparisons test. **f**, Immunoblots and quantification of PAK and CFL activities in the synaptosomes of cultured primary human (n = 3, 3 samples) and mouse cortical neurons (n = 3, 3 samples). *p < 0.05, **p < 0.01, ***p < 0.001; one-way ANOVA with Holm-Sidak’s multiple comparisons test.

**Extended Data Fig. 16.**
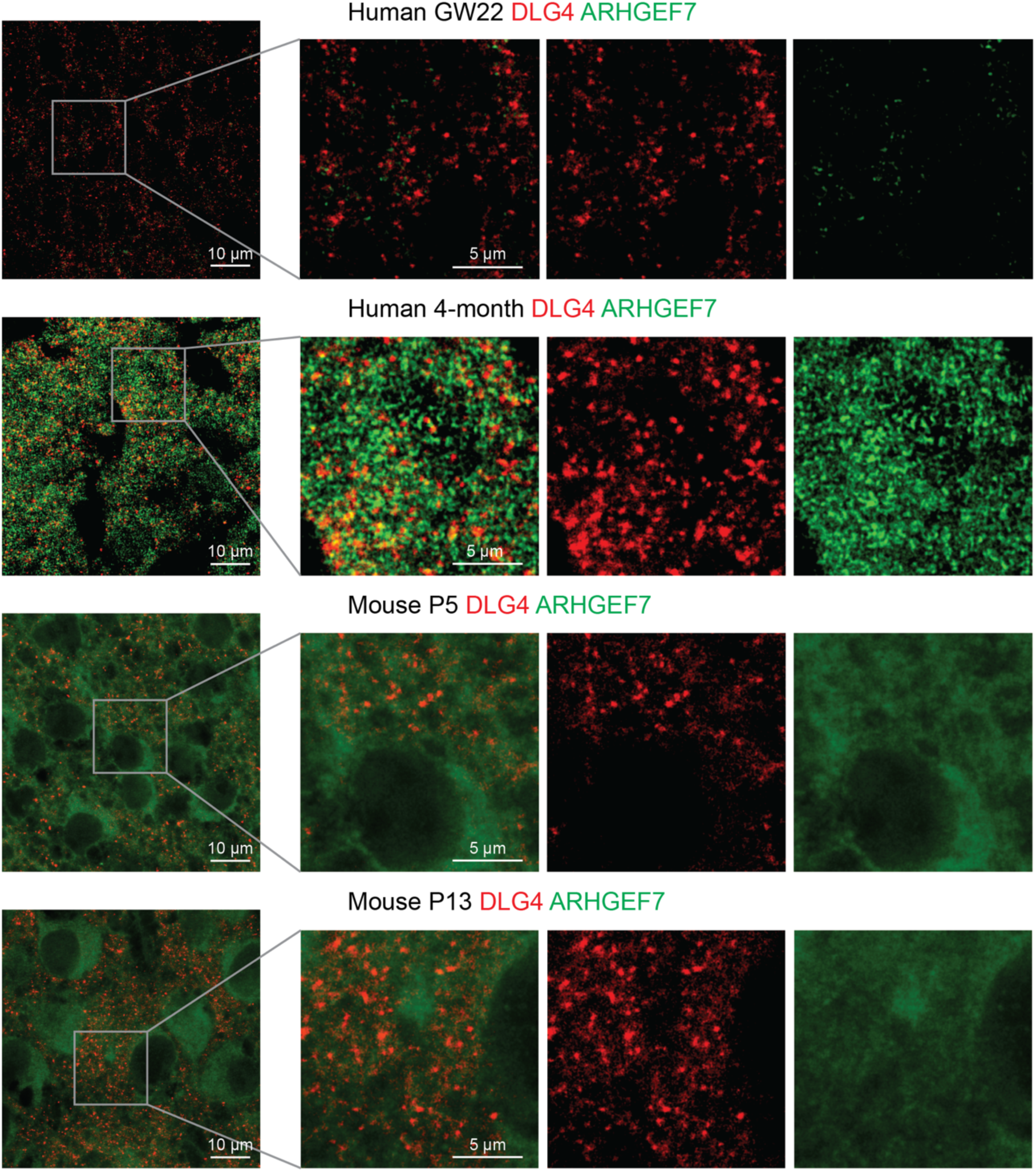
Quantification of ARHGEF7 in the PSD of the human or mouse neocortex. Colocalization of ARHGEF7 with DLG4 in the GW22 human, 4-month human, postnatal day 5 mouse (**c**), and postnatal day 13 mouse neocortex (scale bar: 10 μm or 5 μm as indicated in the figure).

**Extended Data Fig. 17.**
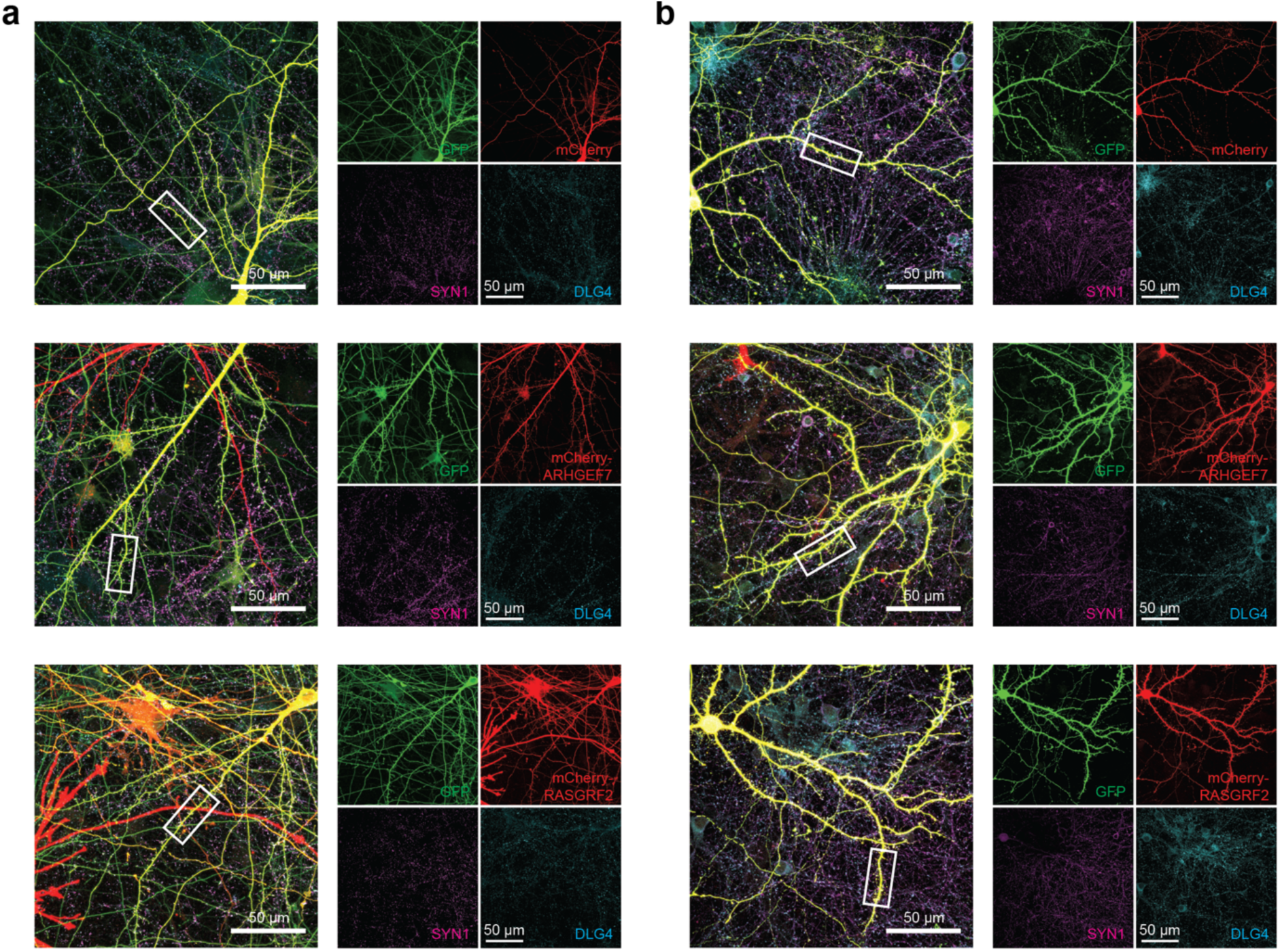
RhoGEF overexpression alters dendritic spine morphogenesis in human and mouse cortical neurons. **a**, Original immunofluorescence images of Fig. 5d. Immunostaining of primary human cortical neurons cultured six weeks *in vitro* transfected with mEGFP-C1 and vectors expressing mCherry, mCherry-ARHGEF7, or mCherry-RASGRF2 (scale bar: 50 μm). **b**, Original immunofluorescence images of Fig. 5e. Immunostaining of primary mouse cortical neurons cultured 8 days *in vitro* transfected with mEGFP-C1 and vectors expressing mCherry, mCherry-ARHGEF7, or mCherry-RASGRF2 (scale bar: 50 μm).

**Extended Data Fig. 18.**
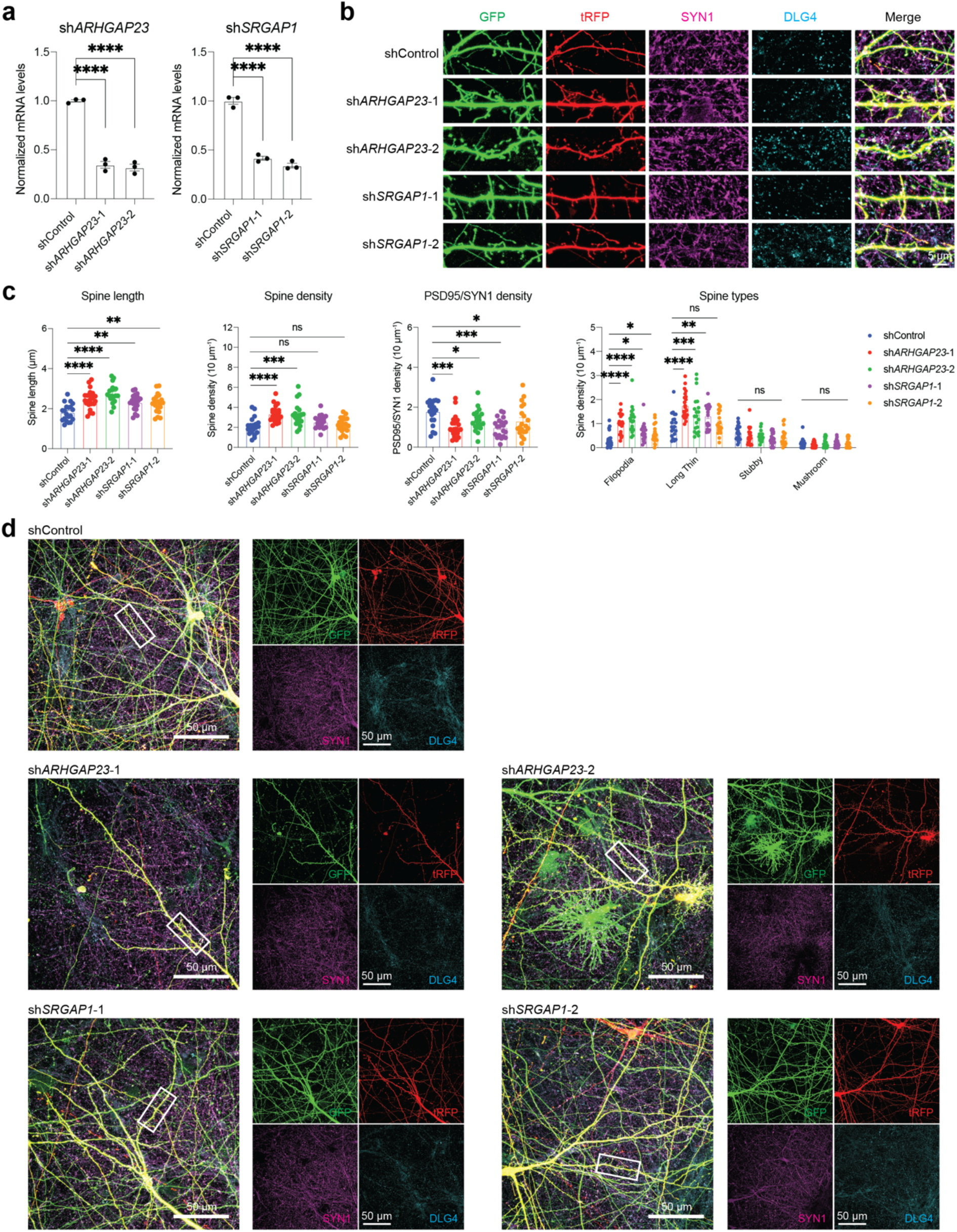
Decrease in selective RhoGAP proteins partially phenocopies RhoGEF overexpression in dendritic spine morphogenesis of human cortical neurons. **a**, Quantification of mRNA levels of *ARHGAP23* and *SRGAP1* in HEK293T cells transfected with control shRNAs (shControl), two shRNAs targeting *ARHGAP23* (shA*RGGAP23*-1 and sh*ARGGAP23*-2), or two shRNAs targeting *SRGAP1* (sh*SRGAP1*-1 and sh*SRGAP1*-2). ****p < 0.0001; one-way ANOVA with Holm-Sidak’s multiple comparisons test. **b**,**c**, Immunostaining of dendrites from primary human cortical neurons cultured six weeks *in vitro* transfected with mEGFP-C1 and vectors co-expressing turbo-RFP (tRGP) and shControl, shA*RGGAP23*-1, shA*RGGAP23*-2, sh*SRGAP1*-1 or sh*SRGAP1*-2 (n = 20, 22, 20, 20, 20 neurons, scale bar: 5 μm). *p < 0.05, **p < 0.01, ***p < 0.001, ****p < 0.0001; one-way ANOVA with Holm-Sidak’s multiple comparisons test. **d**, Original immunofluorescence images of Extended Data Fig. 18b (scale bar: 50 μm).

**Extended Data Fig. 19.**
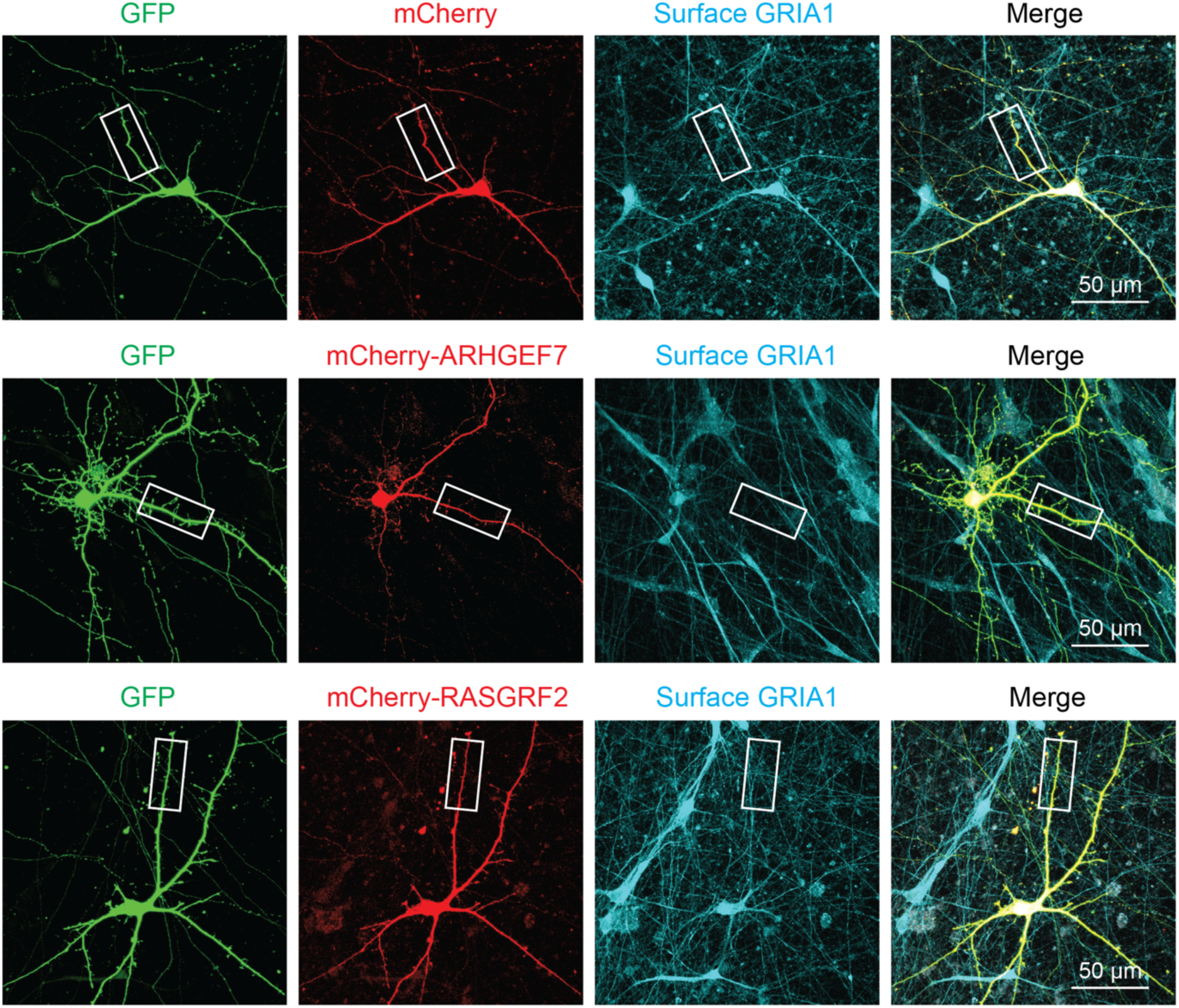
RhoGEF overexpression alters surface α-amino-3-hydroxy-5-methyl-4-isoxazolepropionic acid (AMPA) receptor levels in human neurons. Original immunofluorescence images of Fig. 5g. Immunostaining against surface GRIA1 of primary human cortical neurons cultured six weeks *in vitro* transfected with mEGFP-C1 and vectors expressing mCherry, mCherry-ARHGEF7, or mCherry-RASGRF2 (scale bar: 50 μm).

**Extended Data Fig. 20.**
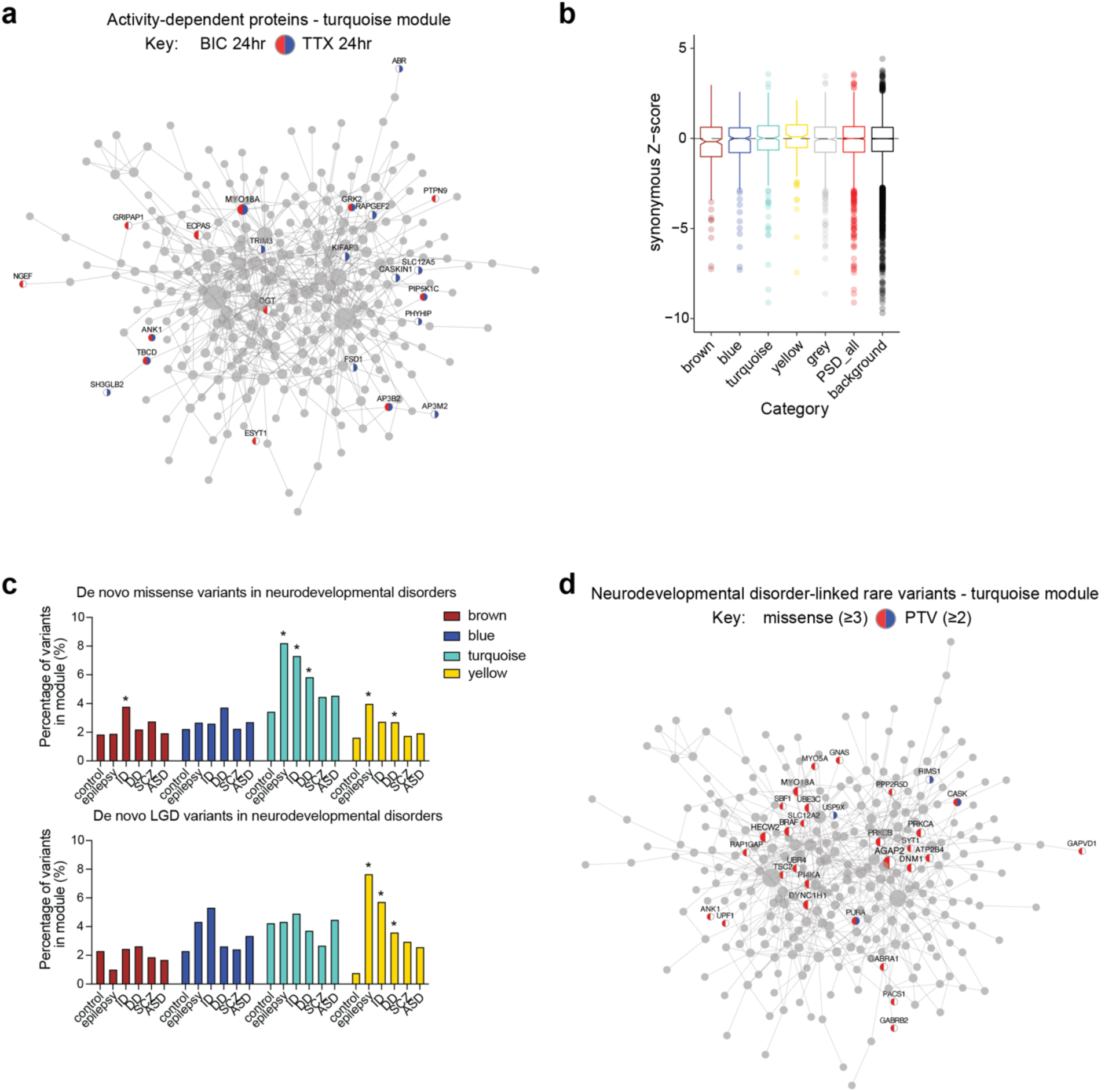
Association of PSD modules with cognitive functions and neurodevelopmental disorders. **a**, PPI-co-abundance network of the turquoise module with activity-dependent proteins highlighted. **b**, Distribution of gnomAD synonymous Z-scores of genes in each category. **c**, Percentage of rare variants located at PSD module genes in subjects with or without neurodevelopmental disorders. The asterisks denote statistically significant differences from the control group (*, p < 0.05; hypergeometric test). **d**, PPI-co-abundance network of the turquoise module with genes carrying neurodevelopmental disorder-linked (at least 3) *de novo* missense variants or (at least 2) PTVs highlighted.

**Extended Data Fig. 21.**
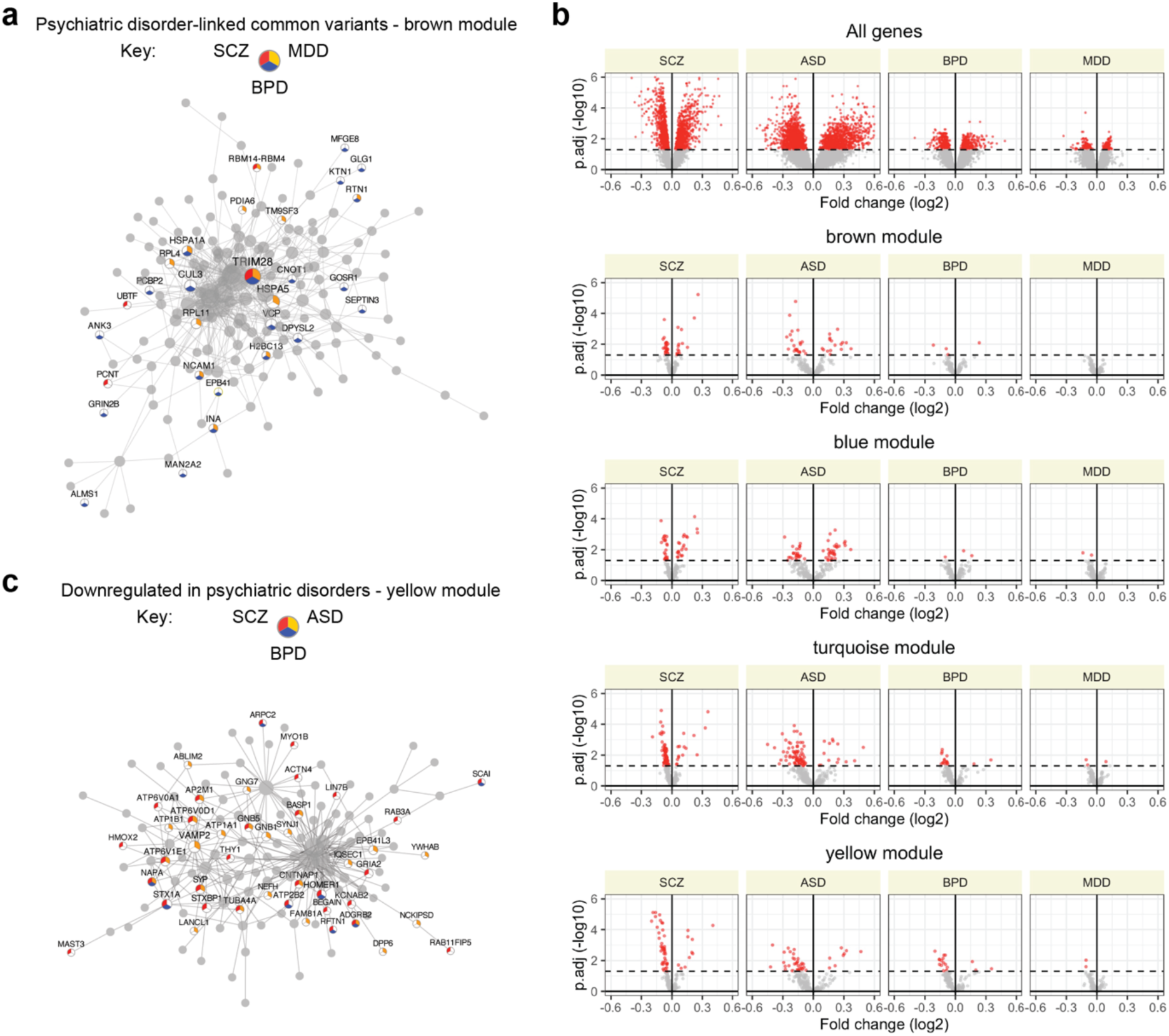
Association of PSD modules with psychiatric disorders. **a**, PPI-co-abundance network of the brown module with genes carrying psychiatric disorder-linked common variants highlighted. **b**, Volcano plots for misexpressed genes after the onset of psychiatric disorders in PSD modules. **c**, PPI-co-abundance network of the yellow module with genes downregulated in psychiatric disorders highlighted.

