## Supplementary figures and images for "A cross-species proteomic map reveals neoteny of human synapse development"

### Supplementary Figures.docx

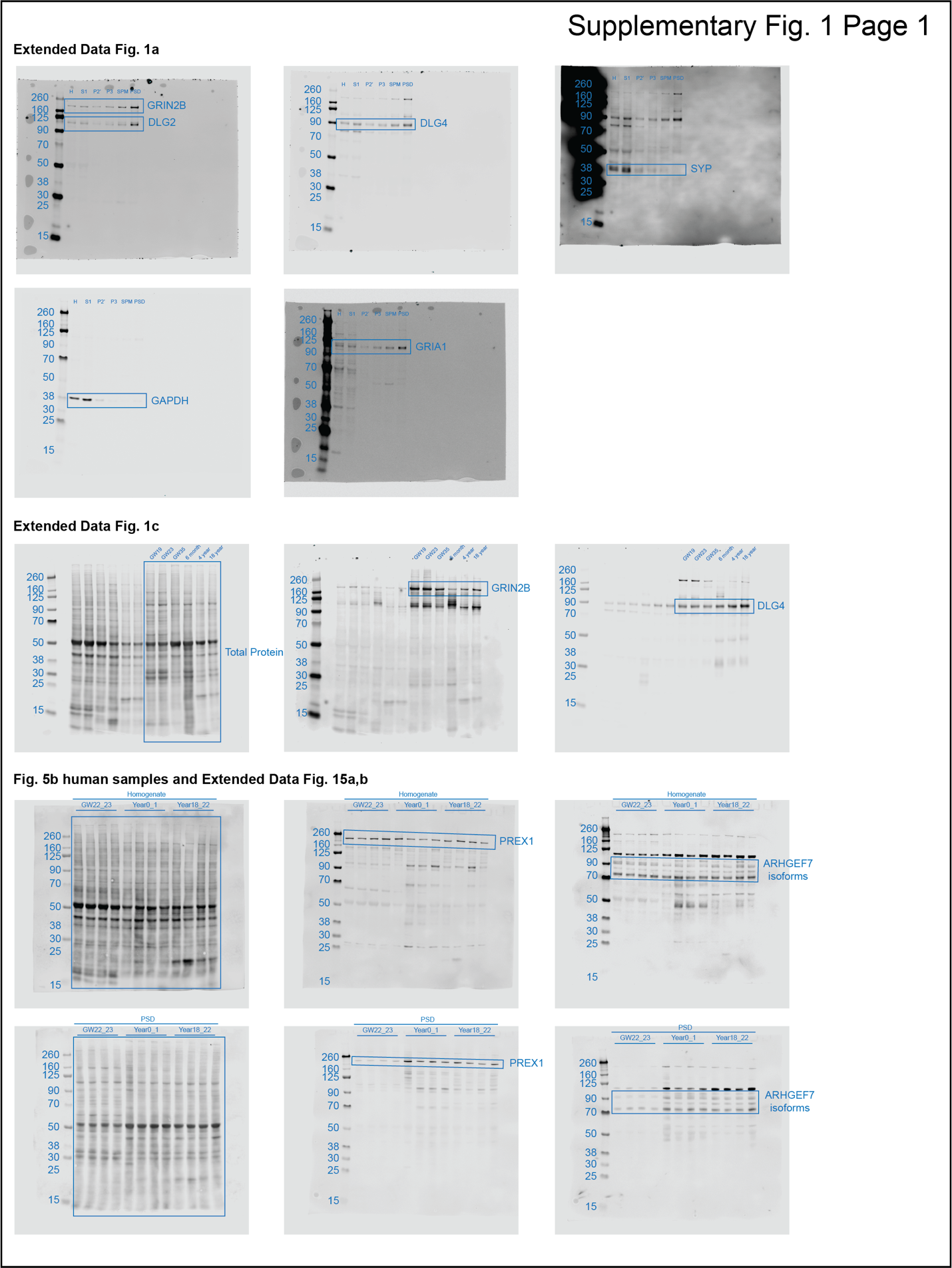

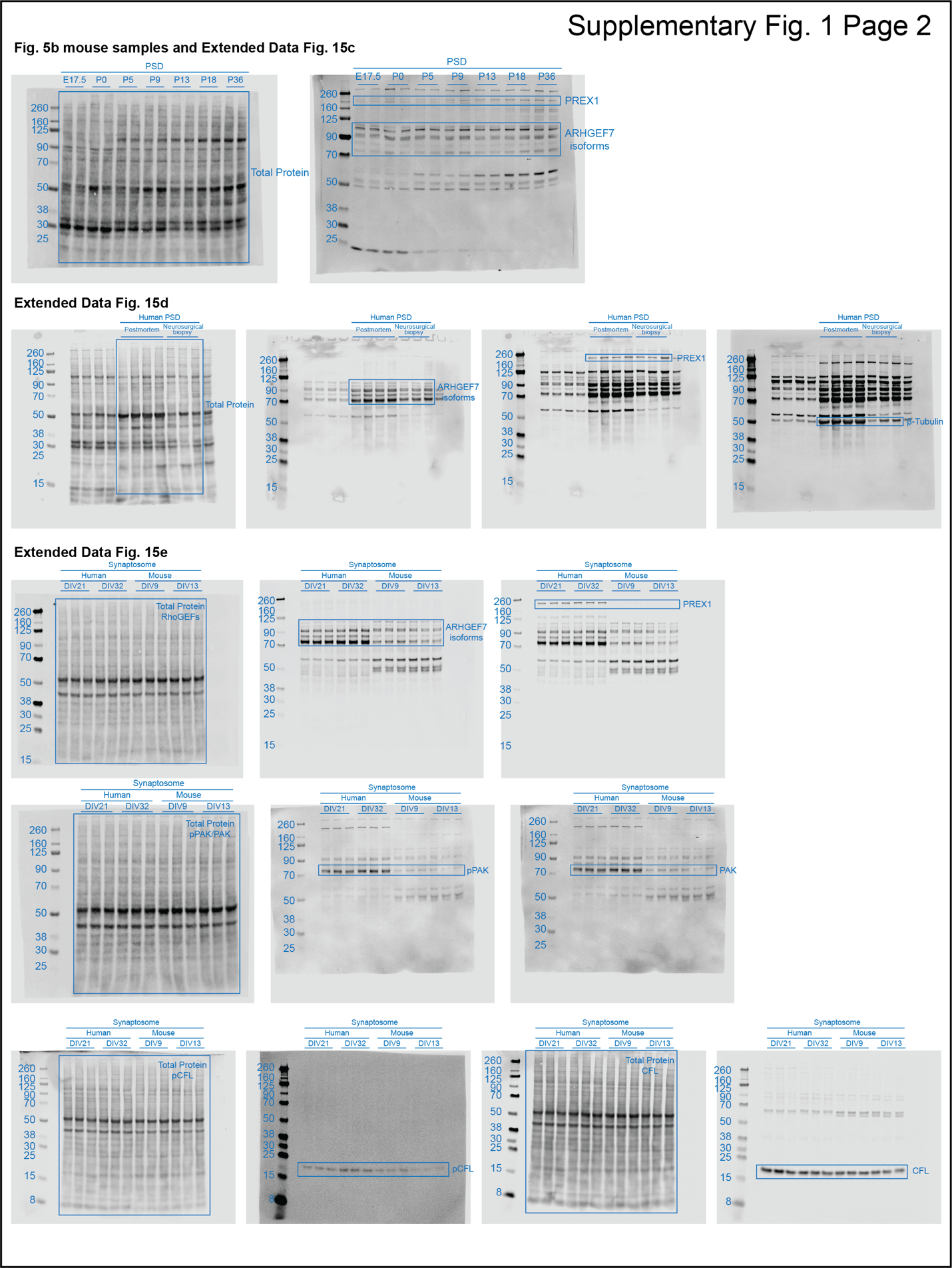

**Supplementary Fig. 1 | Uncropped western blotting images used in this study.**
